# Physical Confinement Modulates the Rate-Limiting Transition in the Release of Phosphate from Actin Filaments

**DOI:** 10.64898/2026.03.12.711388

**Authors:** Kristina M. Herman, Sahithya Sridharan Iyer, Yihang Wang, Thomas D. Pollard, Gregory A. Voth

**Affiliations:** Department of Chemistry, Chicago Center for Theoretical Chemistry, Institute for Biophysical Dynamics, and James Franck Institute, The University of Chicago, Chicago, IL; Department of Molecular Cellular and Developmental Biology, Yale University, New Haven, CT; Department of Molecular Biophysics and Biochemistry, Yale University, New Haven, CT; Department of Cell Biology, Yale University, New Haven, CT

## Abstract

The nucleotide state and rates of transitions between states regulate the dynamics of ATPases. Slow inorganic phosphate (P_i_) release following ATP hydrolysis is often rate-limiting and associated with key conformational changes. Actin filaments offer a unique opportunity to understand the fundamentals of P_i_ release, because identical subunits at filament ends and the interior release P_i_ at markedly different rates. The molecular origin of this difference is debated, so we employed extensive all-atom molecular dynamics simulations to characterize P_i_ release from different subunits within an actin filament. The calculated dissociation rates of P_i_ from ADP-Mg^2+^ in the active site correlate with experimentally measured P_i_ release rates and scale inversely with the numbers of water molecules in the cavity surrounding the γ-phosphate. Simulations show that egress of P_i_ through the protein channels, including through the N111-R177 backdoor, is not rate-limiting and, importantly, that subunits at the filament ends use alternative egress pathways.

**Teaser:** Molecular dynamics simulations show that dissociation of phosphate from Mg^2+^ limits release from all parts of actin filaments

## Introduction

ATP hydrolysis (ATP → ADP-P_i_) and subsequent release of inorganic phosphate (P_i_) (ADP-P_i_ ⇌ ADP) regulate the conformations and dynamics of ATPases. For many ATPases, including myosin, kinesin, and actin, the slow release of P_i_ from the protein is rate-limiting and coupled to conformational changes essential for driving protein function. Thus, P_i_ release serves as the critical regulatory step in the ATPase cycle.

In the case of actin, monomers bind one ATP which is hydrolyzed rapidly and irreversibly upon incorporation of ATP-monomers into a filament.(*1*, *2*) The γ-phosphate (P_i_) dissociates very slowly from internal subunits with a rate constant of 0.002-0.007 s^-1^ (residence time: 2-8 minutes)(*3–8*) producing ADP-actin subunits.(*9*) Note that ADP and ATP exchange into or out of monomers, but not filaments.(*10*) Consequently, phosphate release alone controls aging of actin filaments,(*2*, *11*, *12*) their mechanical properties,(*13*, *14*) rates of assembly and disassembly,(*3*) and affinities for certain actin binding proteins such as cofilin.(*4*, *15*) Thus, phosphate dissociation is the most impactful reaction in the assembly cycle of actin and functions as a clock that marks the local age of the filament.(*11*)

Two independent experimental strategies established that the release of P_i_ from terminal subunits at the ends of filaments is at least two orders of magnitude faster than from interior subunits. Phosphate in the buffer slows the rates of subunit dissociation from both ends of the filaments; calculations constrained by the dependence of these rates on phosphate concentration showed that P_i_ release is at least 600-fold faster (> 2 s^-1^) at both ends than from internal subunits.(*3*) ADP subunits dissociate 27 times faster from barbed ends than ADP-P_i_ subunits (*3*), so single filament imaging of the transition from slow to fast shortening was used to measure the time course of P_i_ release from depolymerizing barbed ends. The estimated rate constant for P_i_ release from barbed end subunits was ∼1.8 s^-1^ (residence time: 0.4 seconds; 260-fold faster than from internal subunits).(*16*) Profilin binding to the barbed end increases the rate of P_i_ release further by two- to five-fold.(*16*)

Bound ligands and certain mutations also influence the rate of P_i_ release. Bound cofilin or coronin increased(*4*, *8*) and bound jasplakinolide decreased(*17*) the rate of phosphate release from internal subunits. The N111S mutant associated with nemaline myopathy(*18*, *19*) increased the rate of P_i_ release ≥15-fold.(*8*)

Major unresolved questions are why the rate of phosphate release from polymerized ADP-P_i_ subunits is so slow, and why the rate varies by location in a filament, with bound ligands, and with mutations? Two hypotheses are proposed. The original idea from steered MD simulations(*20*) of an actin monomer was that the dissociation of P_i_ from the divalent cation (Ca^2+^ in their simulations) is the major energetic barrier for P_i_ release. Subsequent MD simulations of an internal subunit of an ADP-P_i_-actin filament with 5 subunits measured the free energy barrier to break the strong ion-ion interaction between Mg^2+^ and P_i_, which was large enough to make it the rate-limiting step.(*21*) However, this ion pair dissociation hypothesis was not tested to explain the fast rates of release from the filament ends. Other data supported the backdoor gate hypothesis. Short volume-based metadynamics simulations(*8*) identified egress pathways for P_i_ but these authors concluded from a 1.1 μs unbiased simulation that the “backdoor” gate between the sidechains of N111-R177 opens infrequently, and they argued this explains the slow P_i_ release. Cryo-EM structures of filaments generally have closed N111-R177 gates in internal subunits but partially open or open N111-R177 gates in subunits with fast P_i_ release rates (*8*, *22–24*). However, during microsecond-long MD simulations at physiological temperature the N111-R177 gate fluctuates on the nanosecond timescale and is open most of the time, so it would not limit the rate of P_i_ release.(*21*, *25*)

Our objective in this work was to resolve these differences by elucidating the mechanism of P_i_ release from subunits in the interior and at both ends of ADP-P_i_ actin filaments to determine whether the backdoor gate or ion pair dissociation explains the rates of the reaction. We also consider the effect of bound jasplakinolide, a natural product that inhibits P_i_ release.(*17*) Using extensive MD simulations and enhanced sampling methods, we find that the rate of P_i_ dissociation from Mg^2+^ accounts for the relative rates of P_i_ release from the different subunits measured in biochemical experiments. These rates depend on the number of water molecules in the active site, so P_i_ release is slower in interior subunits with few water molecules in the cavity around P_i_ in their active sites than subunits at filament ends with more water molecules and lower barrier heights for dissociation of P_i_ from Mg^2+^. This physical effect is explained by the role of water molecules (and the effective dielectric constant of the environment surrounding the ion pair) in modulating the ion pair stability. Bound jasplakinolide slows P_i_ release by closing the N111-R177 gate and reducing the volume of the phosphate cavity. Furthermore, while subunits in the interior of filaments use the “N111-R177 backdoor” pathway for P_i_ release, the terminal subunits open alternative P_i_ release channels. Our simulations reveal how distinct conformations of subunits at different locations in a filament determine the P_i_ release rates and egress pathways, offering new insights into the mechanisms governing actin filament dynamics and turnover.

## Results

We characterized the dissociation of P_i_ from an actin filament with 13 wild-type subunits and bound ADP-P_i_ based on the cryo-EM structure (PDB 8A2S(*26*)) (Fig. 1A). The large filament size helped to ensure realistic models of the terminal ends and interior subunits of the filament.(*25*) We gradually warmed the cryo-EM structure and subsequently equilibrated the filament at 310 K for 200 ns to allow the filament ends to relax before production runs totaling 1.6 μs. Past work showed that ∼200 ns was necessary to equilibrate the ends of the filament during simulations at 310 K.(*12*) In addition, we equilibrated filaments with 13 ADP-P_i_ actin subunits with bound jasplakinolide (Fig. 1D), a natural product that inhibits P_i_ release.(*17*)

**Figure 1.**
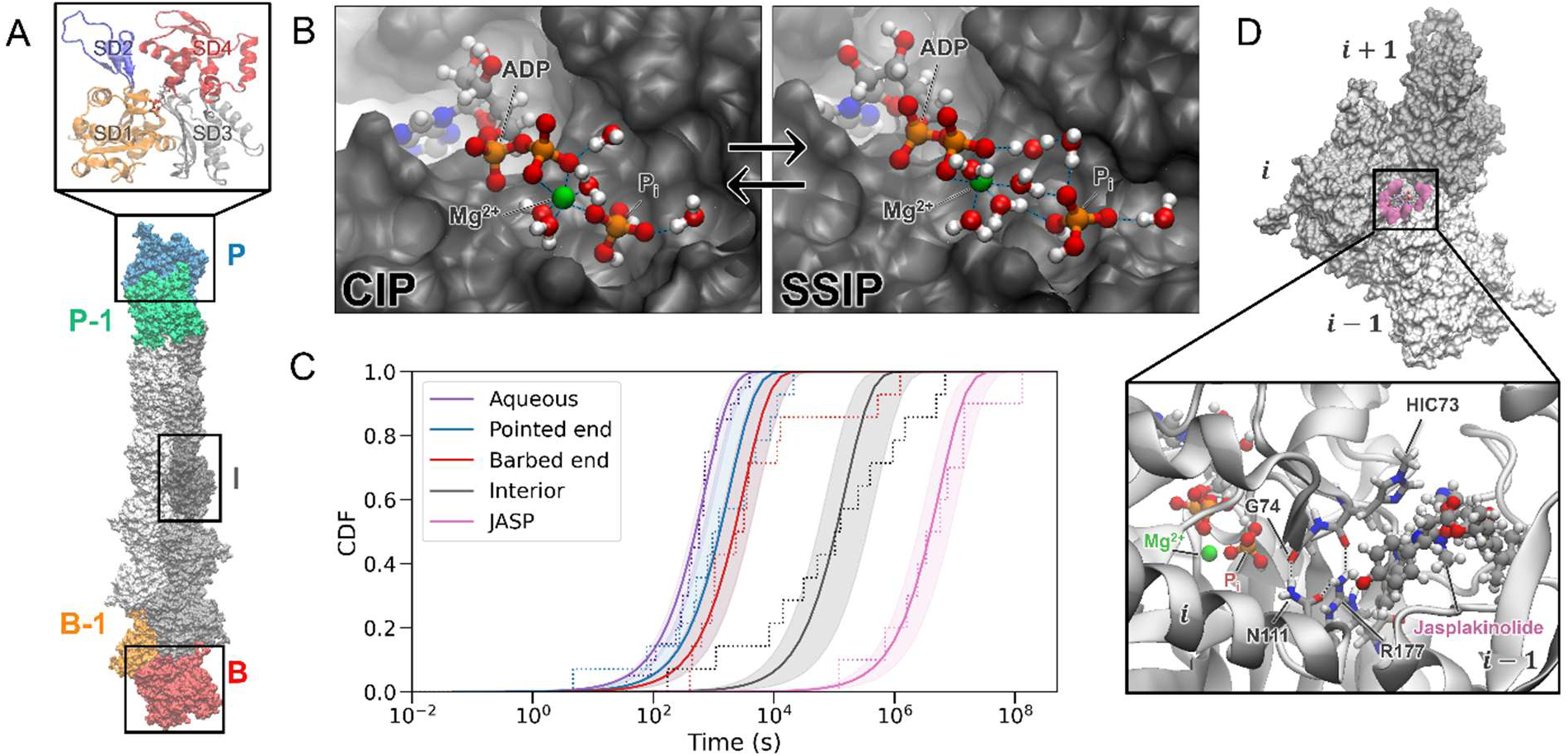
The transition from the CIP state to the SSIP state in actin subunits during a simulation of a filament with 13 ADP-P_i_-actin subunits. (A) Space-filling model of an actin filament with 13 subunits after 200 ns of equilibration [interior subunits: gray, terminal pointed end subunit (P): blue, penultimate subunit (P-1): green, terminal barbed end subunit (B): red, penultimate subunit (B-1): orange]. The zoomed inset shows a ribbon diagram of an actin subunit with ADP-Mg^2+^-P_i_ in the active site. Different colors distinguish the 4 subdomains of the protein. (B) Snapshots of the active site of an interior subunit in the CIP state (left) and SSIP (right) state with CPK models of ADP, P_i_, Mg^2+^ and all water molecules with oxygen atoms within 4.5 Å of the phosphorus atom of P_i_ or within 2.5 Å of the magnesium atom. The protein surface in dark gray was computed in VMD(33) with a probe radius of 1.4 Å. (C) The empirical cumulative distribution functions (CDFs, dotted lines) of the transitions from CIP to SSIP in TIP3P water (purple), the pointed end subunit (blue), the barbed end subunit (red), the interior subunit (dark gray), and an interior subunit with jasplakinolide bound (pink). The stairstep dotted lines are the time points where each successive fraction of the simulations transitioned. The solid lines show the fit to 1 − *e*^*k*_0_*t*^ where *k*_0_ is the rate constant. The shaded regions reflect the uncertainty obtained by bootstrapping. (D) Space-filling model with jasplakinolide bound at the interface of three neighboring actin subunits (*i*+1, *i*, *i*-1). Surfaces of the subunits within 5 Å of jasplakinolide are colored pink. The zoomed inset shows jasplakinolide interacting closely with the sensor loop and R177-N111 gate of subunit *i*. The dotted lines highlight hydrogen bond interactions between G74-N111, N111-R177, and R177-HIC73.

### The rates of transition from contact ion pair (CIP) to solvent-separated ion pair (SSIP) in subunits of actin filaments

Figure 1B shows an internal actin subunit with ADP-Mg^2+^-P_i_ in the nucleotide binding cleft in the contact ion pair (CIP) state with direct interactions between P_i_ and Mg^2+^. Note the few water molecules in the first solvation shell of P_i_ in this restricted space. Cryo-EM structures of ADP-P_i_ actin filaments (PDB 8A2S,(*26*) 6DJN,(*22*) 8D14(*14*)) exist in the CIP state. To dissociate from the subunit, P_i_ must first transition from CIP (ADP-Mg^2+^-P_i_) to SSIP (ADP-Mg^2+…^P_i_) (Fig. 1B). It is well-established that water molecules play a critical role in stabilizing and facilitating the CIP-to-SSIP transition.(*27–30*) Mechanistically, the hydrogen bond network, including interactions between Mg^2+^, P_i_ and water, must rearrange to separate the ion pair as illustrated in Fig. 1B (right).

We compared the rates of transition from CIP to SSIP for subunits in the middle and the ends of filaments to learn whether this transition accounts for the differences in P_i_ dissociation rates in biochemical experiments. Knowing from biochemical experiments that P_i_ release is very slow, we used well-tempered metadynamics (WT-MetaD), an enhanced sampling method, to simulate the CIP-to-SSIP transition in filaments with 13 ADP-P_i_-subunits. WT-MetaD(*31*, *32*) accelerates sampling along select degrees of freedom—termed collective variables (CVs)—that capture the slow, relevant motions underlying the transition of interest. During the MD simulations, biases are deposited along the CVs to accelerate the sampling of these degrees of freedom, enabling the exploration of “rare events”, such as the CIP-to-SSIP transition, within timescales accessible by simulation.

Naturally, the accelerated sampling along the selected CVs changes the observed kinetics of the ion pair dissociation. Using the method of Tiwary and Parrinello,(*34*) we estimated the time-acceleration factor *α* as the time integral of the bias (see Methods for details), which was deposited along the distance between Mg^2+^ and the center-of-mass of P_i_, *r* (P_i_-Mg^2+^), during MD simulations. Each simulation was initiated from the CIP state and ended once the system transitioned to the SSIP state, yielding a rescaled passage time for the CIP-to-SSIP transition. Importantly, each independent simulation started from a different initial structure that was sampled every 20 ns from an unbiased simulation. This helped to produce statistics representative of the ensemble of conformations at 310 K(*25*) by sampling across protein fluctuations on the ns to μs timescale. We note that this method is more accurate for estimating a rate than the application of Eyring theory (transition state theory).

We calculated the empirical cumulative distribution functions (CDF) from rescaled CIP-to-SSIP transition times for five different actin subunits and for ADP-Mg^2+^-P_i_ in water (Supplemental information sections 1 and 2). The CDFs (dotted lines in Fig. 1C) are the fraction of simulations that transitioned to SSIP at time *t* on the *x*-axis. Fitting the CDFs (to *CDF* = 1 − *e^-k_0_t^*) yielded estimates of the rate constant *k*_O_ for each system. The shaded regions in Fig. 1C represent the uncertainty in the rate constant estimated by bootstrapping. The CIP-to-SSIP transition for ADP-Mg^2+^-P_i_ is faster in water (purple in Fig. 1C) than in any of the actin subunits.

We relied on the Kolmogorov-Smirnov (KS) test for statistical evaluations regarding whether the observed transition times come from the fitted distributions. The results for all the systems pass the KS test (*p* > 0.05) with the lowest *p* value being 0.17 (Table 1), indicating that the calculated transition times come from the fitted Poisson distributions (solid lines, Fig. 1C).

**Table 1.** Summary of the calculations of the rate constants for the transition from CIP to SSIP (*kCIP*→*SSIP*) for six systems. The table also lists the number of independent CIP-to-SSIP transition times (*n*), *p* values from the KS test, rates relative to internal subunits, and estimated differences in barrier heights relative to internal subunits (*ΔΔG^‡^interior*). The values in brackets indicate the uncertainties obtained from the standard error via bootstrapping.

|  | Aqueous | Pointed End | Barbed End | Interior | JASP-bound |
| --- | --- | --- | --- | --- | --- |
| Number of simulations | 20 | 14 | 14 | 14 | 10 |
| $k_{CIP \rightarrow SSIP}$ ($10^{-6} \text{ s}^{-1}$) | 1400 | 540 | 310 | 6.5 | 0.18 |
| [range] | [930 – 1700] | [180 – 1000] | [180 – 600] | [2.0 – 13] | [0.084 – 0.29] |
| $p$ value | 0.85 | 0.28 | 0.26 | 0.17 | 0.68 |
| Simulated rate relative to interior subunit [range] | 215x<br>[72 - 850x] | 83x<br>[14 - 500x] | 48x<br>[14 - 300x] | 1 | 0.028x<br>[0.14 - 0.006x] |
| $\Delta\Delta G^\ddagger$ (kcal/mol) estimated from rate | 2.6 - 4.2<br>lower | 1.6 - 3.8<br>lower | 1.6 - 3.5<br>lower | 0 | 1.2 - 3.1<br>higher |

We note that this rate calculation computed from rescaled first passage times using 1 CV, *r* (P_i_-Mg^2+^), yields the same rate constant as transition state theory using the free energy barrier (obtained with 2 CVs) for ADP-Mg^2+^-P_i_ in water (Section 1 and 2 of SI). This agreement shows that the 2^nd^ CV describing coordination of Mg^2+^ is not necessary to *promote* the transition of CIP to SSIP, although it is necessary to separate the CIP, SSIP, and transition states in the PMF.

The CHARMM36m potential used for the simulations overstabilizes the CIP state (see Section 1 in SI for benchmarking of force fields). The overstabilization of the CIP (and the exponential relationship between the barrier height and rate) yields CIP-to-SSIP transition rates slower than experimental measurements of P_i_ release. For this reason, Table 1 also lists the rates relative to the interior subunits, enabling a relative comparison of how the barrier heights differed in the sampled subunits.

The rates of the transitions from CIP to SSIP calculated from the simulations are faster for the terminal subunits at both ends than interior subunits: ∼ 48-fold faster in the barbed end subunit and ∼ 83-fold faster in the pointed end subunit (Table 1). These relative rates differ only ∼ 6-fold from biochemical measurements which used indirect measurements of the phosphate release rate from the subunits at the filament ends.(*3*, *7*)

The natural product jasplakinolide binds between three subunits in actin filaments (Fig. 1D) and inhibits phosphate release from ADP-P_i_-actin filaments in experiments, though the rate was not measured.(*17*) Bound jasplakinolide slows the CIP-to-SSIP transition 36-fold in the simulations of interior subunits (Fig. 1C, Table 1).

We simulated the CIP-to-SSIP transition for the N111S mutant. SI section 7 explains that the rate was similar to WT as expected from the small differences in the overall rate of phosphate release. The rate constants calculated for the transition from CIP-to-SSIP ranged 7,500-fold (ΔΔ*G*^‡^ = 5.4 kcal/mol) from slow transitions at 10^-3^ s^-1^ for ADP-Mg^2+^-P_i_ in water to very slow transitions at 10^-7^ s^-1^ in interior subunits with bound jasplakinolide.

Most importantly, the relative rates of the CIP-to-SSIP transitions agree with the relative rates and the rank order of the rates of phosphate release from actin in biochemical experiments. The differences of the calculated relative rates to the experimental ones equate to about 1.3 kcal/mol difference in energy barrier, though they agree closely with experiments within uncertainty. Moreover, the CIP-to-SSIP transition is but one key part of the P_i_ dissociation process (though the main one). Other features along a given egress pathway will further affect the rate of P_i_ release rates, though we estimate these to be marginal (see later text). The general agreement of these numbers supports the assignment of the CIP-to-SSIP transition as the rate-limiting step in P_i_ release from different parts of filaments.(*21*)

### The physical origin of the different rates observed for the CIP-to-SSIP transition

Since water molecules facilitate the CIP-to-SSIP transition of simple ions in liquid water(*27–30*), we examined the water molecules in the active sites to explore the origin of the range of calculated rates for the CIP-to-SSIP transitions in the different parts of actin filaments (Fig. 1C). We defined the volume available to water within 5 Å of the phosphorus atom of P_i_, while excluding the volume occupied by protein, Mg^2+^, and ADP (Fig. 2A, B). We computed these “phosphate cavities” (Fig. 2A) across unbiased MD simulations with POVME3(*35*).

**Figure 2.**
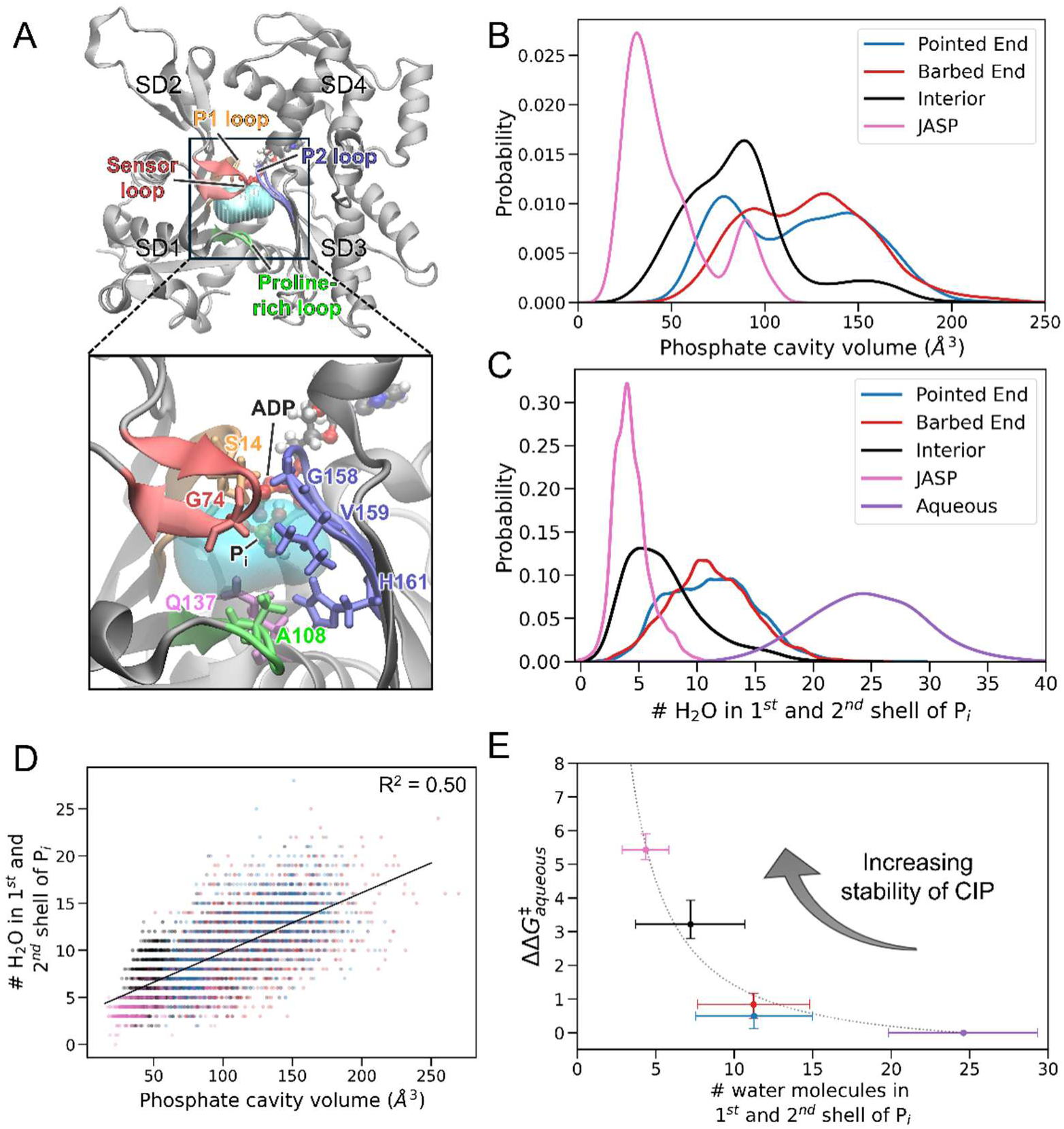
Relation of water molecules in the phosphate cavity and the stability of CIP. (A) Ribbon diagram of an internal subunit in an ADP-Mg^2+^-P_i_-actin filament with the phosphate cavity volume in cyan delimited by protein residues. The phosphate cavity volume is defined the volume within 5 Å of the phosphorus atom of P_i_ excluding contributions from Mg^2+^, ADP, and the protein. Amino acids in the P1 (orange), P2 (blue), sensor (red), and proline-rich (green) loops are labeled in the top panel and colored. Some of their sidechains near P_i_ are shown in licorice representation. (B) The phosphate cavity volumes and (C) the number of water molecules in the 1^st^ and 2^nd^ solvation shells of P_i_ in four actin subunits: pointed end, barbed end, interior, and interior with jasplakinolide bound. (D) The frame-to-frame correlation between the phosphate cavity volume and the number of water molecules in the 1^st^ and 2^nd^ solvation shell of P_i_. The solid black line denotes the line of best fit. (E) Correlation between the number of water molecules near P_i_ and the estimated barrier height of the CIP-to-SSIP transition relative to the dissociation in water (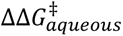, kcal/mol). The relative barrier heights on the *y*-axis are derived from the computed rates of CIP-to-SSIP transition using the Eyring equation. The vertical error bars represent the uncertainties in the barrier height (derived from the uncertainty in the computed rates). The horizontal error bars show the standard deviation of the number of water molecules in the first and second shells around P_i_ over a simulation. The dotted line shows the line of best fit 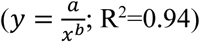 and is primarily used to guide the eye.

Owing to rapid fluctuations of the conformations of the subunits,(*25*) the volumes of these phosphate cavities varied over time and differed for the five subunits considered (Fig. 2B). The volumes of the phosphate cavities (Fig. 2B) and the numbers of water molecules in the 1^st^ and 2^nd^ solvation shells of P_i_ (Fig. 2C) were smallest for interior subunits in filaments with bound jasplakinolide, followed by interior subunits, and the subunits at both ends. The purple curve in Fig. 2C shows the water molecules in the 1^st^ and 2^nd^ solvation shells of P_i_ during a simulation of ADP-Mg^2+^-P_i_ in water, highlighting the substantial reduction of water molecules near P_i_ in the phosphate cavities. The means of the distributions varied by a factor of ∼ 3 for the actin subunits considered (Fig. 2C).

The number of water molecules near P_i_ is proportional to the volumes of the five phosphate cavities we characterized (Fig. 2D). The cavity volumes and the number of water molecules are related yet distinct, because POVME3 reports the volume of regions available to water but water molecules occupying the phosphate cavity differ depending on their shapes and the amino acids near P_i_.

The P_i_ anion preferred slightly different positions in the phosphate cavities of subunits across the filament (Table S1). In interior subunits, P_i_ interacted closely with the top of the phosphate clamp (P1 loop, residues 13-14; and P2 loop, residues 154-159). In subunit P, P_i_ favored a position deeper in the phosphate cavity, at the base of the P1 loop and near the proline-rich loop. In subunit B, P_i_ favored a more forward position in the phosphate cavity, at the base of the P1 loop.

To understand the role of the protein cavity in modulating the barrier heights, we used the Eyring equation (transition state theory) and the rate constants computed in Fig. 1C to estimate barrier heights for the transitions from CIP to SSIP relative to the transition in water (Fig. 2E).

Strikingly, these barrier heights for the CIP-to-SSIP transition (relative to water) scaled inversely with the number of waters surrounding P_i_ (Fig. 2E). Note that the *x*-error bars reflect the standard deviation (rather than uncertainty in the mean) of the distribution of the number of water molecules near P_i_ (Fig. 2C) over 1.6 μs of unbiased simulation. This relationship indicates that large volumes with more water molecules (higher effective dielectric screening) favor the transition from CIP to SSIP. Conversely, the ion pairing is strong in subunits of filaments with bound jasplakinolide which have small phosphate cavities and numbers of water molecules, slowing the transition from CIP to SSIP.

### Relationship between subunit conformations and phosphate cavity volume

The dynamic fluctuations of the subunit conformations in actin filaments at 310 K shape the distributions of phosphate cavity volumes in the subunits where amino acids near Mg^2+^-Pi can exclude water molecules from around the ion pair.

Thermal motions cause the conformations of subunits in actin filaments to fluctuate on a nanosecond time scale(*25*) about two major normal modes:(*36*) a propeller twist of the dihedral angle between SD1/2 and SD3/4 and an orthogonal scissors motion that separates SD2 and SD4 (Fig. 3A). Both motions are hinged below the bound nucleotide, so they influence the volume of the phosphate cavity and numbers of water molecules near P_i_.

**Figure 3.**
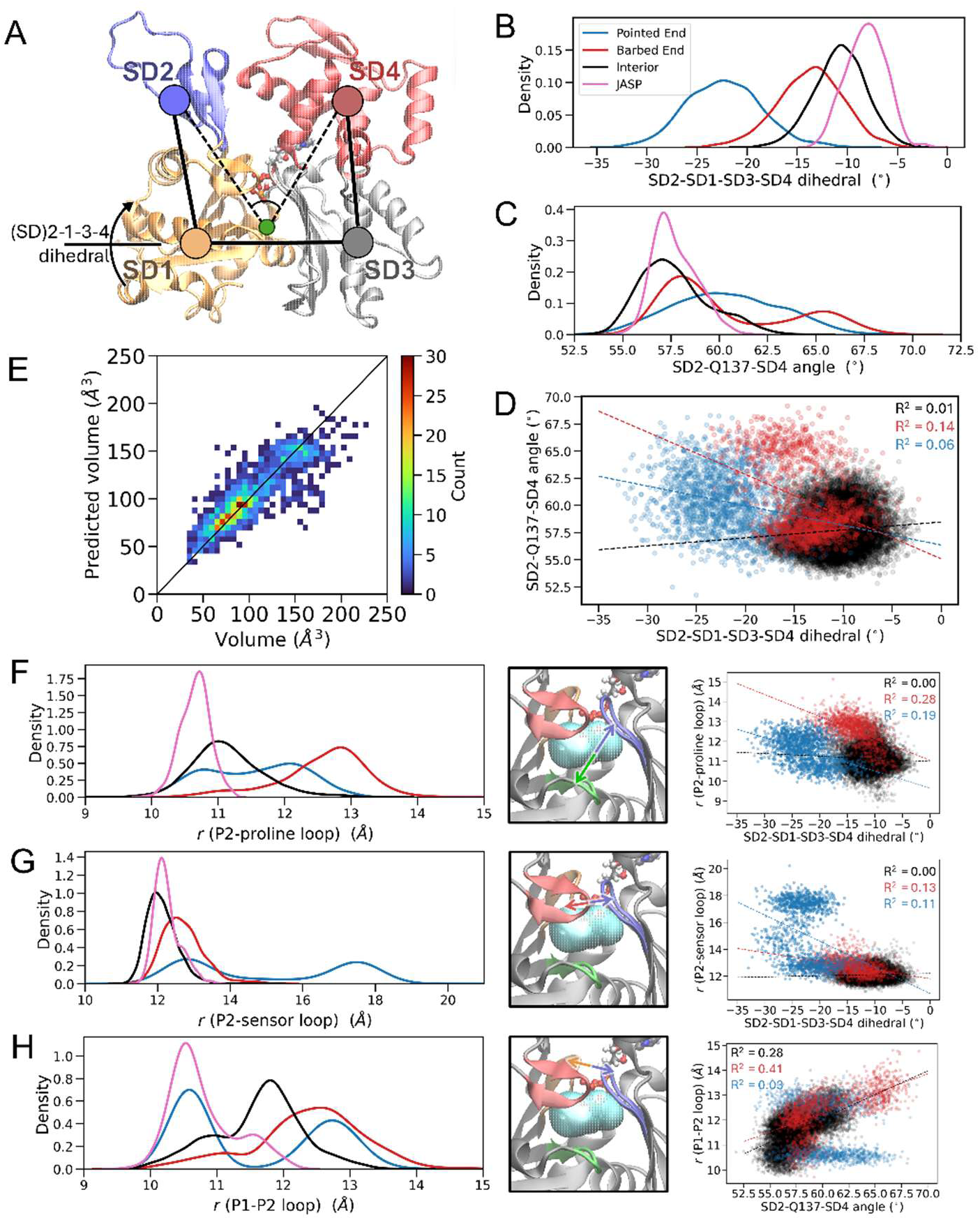
Actin subunit conformations in filaments and their relationship to the phosphate cavity volume. (A) Actin subunit defining the centers-of-mass of the four subdomains, the scissors angle (SD2-Q137-SD4), and the SD2-SD1-SD3-SD4 dihedral angle. (B-C) Distributions of the (B) SD2-SD1-SD3-SD4 dihedral angles and (C) scissors angles (SD2-Q137-SD4) in 5 actin subunits during unbiased MD simulations; (D) Frame-to-frame correlation between the scissors (SD2-Q137-SD4) and SD2-SD1-SD3-SD4 dihedral angles plotted at an interval of 1 ns across ∼1.6 μs of unbiased simulation time; (E) Correlation between volumes predicted by a Random Forest Regression model on the *y*-axis and volumes of the phosphate cavity on the *x*-axis at each frame of the test set of ∼ 1,300 frames. We trained a Random Forest Regressor (*y*-axis) using the COM-to-COM distances between the P1, P2, sensor, and proline-rich loops along with the scissors and dihedral angles from ∼6,300 MD frames. The mean absolute error (MAE) of the model on the test set (∼1,200 frames) is 14 Å^3^; (F-H) Distributions of (F) P2 – proline-rich loop distances; (G) P2 – sensor loop distances; and (H) P1-P2 loop distances in four actin subunits during unbiased MD simulations. The statistics for interior subunits were aggregated across B-2 to P-2 subunits. The middle panels of D-F are ribbon models of the interior subunit with the volume of the phosphate cavity in cyan and colored arrows to highlight the two loops used to compute their center-of-mass to center-of-mass distance. The centers-of-mass were computed using the C_α_ atoms of residues 11-16 (P1loop, orange), 71-77 (sensor loop, red), 107-111 (proline-rich loop, green), and 154-161 (P2 loop, blue). The right panels of D-F show the relationships between the relative loop positions and the normal mode motions plotted at an interval of 1 ns. The dashed lines show the lines of best fit for each subunit.

Variations in time of the dihedral and scissors angles are weakly correlated in our simulations of subunits at the filament ends and uncorrelated in interior subunit (Fig. 3D), so these normal modes largely fluctuate independently. These intramolecular fluctuations are coupled with the relative positions of the P1, P2, sensor, and proline-rich loops which border the phosphate cavity (right panels, Fig. 3F-H). Correlations between the relative positions of the P1, P2, sensor, and proline-rich loops and the phosphate cavity volume are weak to moderate (Fig. S13), but no single structural parameter *alone* fully accounts for the volume of the phosphate cavity. This section first examines each of those parameters and then shows how they collectively determine the volume of the phosphate cavity.

Each subunit analyzed had different distributions of dihedral and scissors angles (Figs. 3B, C) and different distributions of residues close to the heavy atoms of P_i_ (Table S1). The wide distributions of the dihedral and scissors angles for twisted subunit P show that the free energy surface along these degrees of freedom is flatter than for other subunits (Figs. 3B, C),(*25*) allowing broad sampling of these degrees of freedom within the timescales of our simulations. At the barbed end, the distributions of dihedral angles (Fig. 3B) and scissors angles (Fig. 3C) overlap those of interior subunits, but the distribution of dihedral angles is broader and shifted ∼3° to more twisted angles and the distribution of scissors angles has an additional small peak at about 66°. The latter may be related to the propensity for SD4 of subunit B to lose contacts with SD3 of subunit B-2 in simulations.(*12*, *25*) Altogether, the difference between the major normal mode fluctuations of interior and terminal subunits highlights the importance of lateral and longitudinal interactions in stabilizing the flattened subunits with smaller SD2-SD4 distances at the filament interior.

Two internal distances are weakly correlated with fluctuations of the dihedral angle: the distances between the P2 loop in the inner domain and both the proline-rich and the sensor loops in the outer domain (Fig. 3F-G). Consequently, these distances are larger in more twisted internal subunits. Oosterheert et al. identified this motion as contributing to the relocation of water molecules in the phosphate cavity during polymerization of an actin subunit.(*26*) The barbed end subunit has the largest distances between the P2 and proline-rich loops, which may result from having free subdomains 1 and 3 at the end of the filament. The proline-rich loop of subunit B was also repositioned in a cryo-EM structure.(*8*)

The scissors motion influences the distance between the P1 and P2 loops (Fig. 3H). Subunits at the filament ends exhibit larger angles and a greater range of fluctuations than interior subunits.

The direct, frame-to-frame correlations between the normal mode motions and the volume of the phosphate cavity are generally weak (Fig. S12). The correlations are strongest in subunit B with R^2^ = 0.19 for the dihedral angle and R^2^ = 0.38 for the scissor angle.

While many of parameters correlate weakly with the volume of the phosphate cavity (Fig. S12-S13), collectively the normal mode motions together with the relative positions of the four loops bordering the phosphate cavity determine the cavity volume (Fig. 3E). We produced a dataset of these variables over the MD simulations to train a Machine Learning-based Random Forest Regressor(*37*) to predict the phosphate cavity volume. We used ∼3,100 frames from interior subunits and ∼3,100 frames from the filament ends. The trained model yielded a mean absolute error of 14 Å^3^ and a mean absolute percent error of 15% on a test set of ∼1,300 MD frames (Fig. 3E). This minimal, interpretable model focused on backbone fluctuations in the subunit. Fig. S14 shows a higher accuracy model (MAE on test set: 10 Å^3^) that includes details about the side chain positions of residues in the phosphate cavity.

To summarize, the relative positions of the loop regions are the strongest predictors of the phosphate cavity volume (Fig. S13, S15), and their fluctuations are driven by the normal mode motions. The subunits at the end of the filament (which lack certain longitudinal and lateral interactions) exhibit larger fluctuations in the dihedral and scissors angles (Fig. 3B-C) and, consequently, larger distances between the loops which border the phosphate cavity (Fig. 3F-H).

### Effects of bound jasplakinolide on phosphate cavity volumes of interior subunits

A cation-π interaction between R177 and jasplakinolide reinforces the N111-R177 gate, stabilizes the sensor loop, and interacts with residues (190–201) in SD4 of the lateral neighbor *i* − 1 subunit (Fig. 1D). The smaller phosphate cavities in subunits with bound jasplakinolide (Fig. 2A) result from the stabilized scissors angle (Fig. 3C) and flattened dihedral angles (Fig. 3B, mean -8°). Owing to constrained sidechain arrangements in the active site, fluctuations in the phosphate cavity volumes correlate more strongly than in wild type filaments with fluctuations in the dihedral angle (R^2^ = 0.39), the scissors angle (R^2^ = 0.39) and the P1-P2 loop distance (R^2^ = 0.70) in simulations with bound jasplakinolide (Fig. S16).

### Phosphate egress pathways from subunits across the filament

WT-MetaD simulations, biasing only the distance between Mg^2+^ and the center-of-mass of P_i_, revealed the pathways used to release P_i_ from the pointed end, interior, B-2, B-1, and barbed end subunits (Fig. 4). This collective variable is inherently agnostic to the egress pathway. Depositing small, gentle biases (width: 0.05 Å) every 10 ps allowed P_i_ to explore the active site and the possible pathways, resulting in physically plausible pathways. We aimed to identify the *most favorable* egress pathway(s) rather than sample diverse pathways.

**Figure 4.**
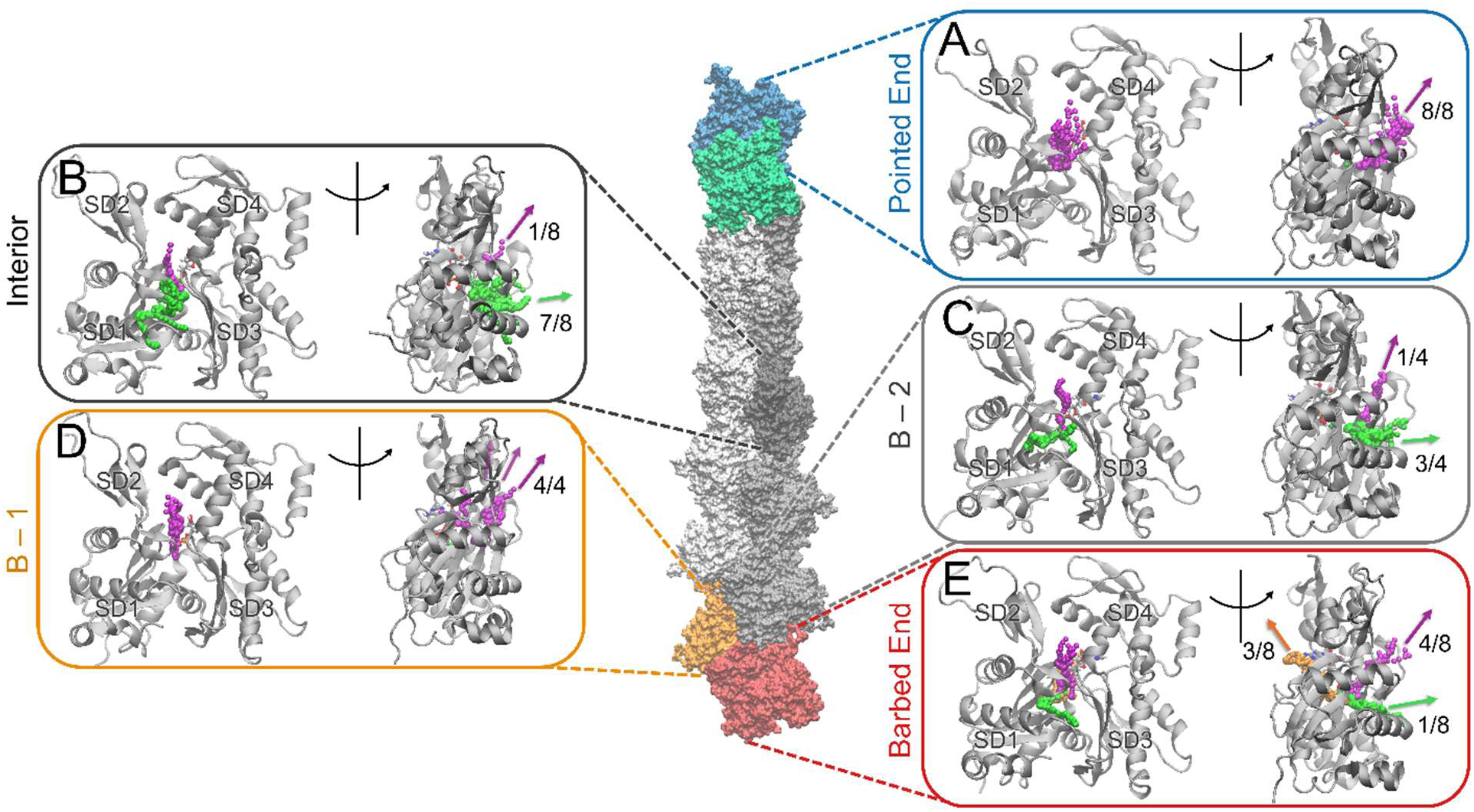
Egress pathways from five subunits during enhanced sampling simulations of the filament with 13 ADP-P_i_ subunits: (A) pointed end subunit P; (B) interior subunit I; (C) barbed end subunit B-2; (D) barbed end penultimate subunit B-1; and (E) barbed end terminal subunit B. Eight independent trials were simulated for the pointed end, interior, and barbed end subunits. Four independent trials were simulated for the B-2 and B-1 subunits. The colored beads represent the positions of P_i_ at different snapshots in time. The colors of the beads and arrows highlight 3 categories of egress pathways: purple, back door above sensor loop; green, back door below sensor loop; and orange, front door.

The deposited bias can be used to estimate the PMFs (Fig. S23) and timescales (Fig. S24) for egress from the SSIP state to the protein exterior. These timescales typically range from the microsecond to millisecond; much faster than the slow dissociation of P_i_ from Mg^2+^ in the phosphate cavity. These estimated rates should be considered lower bounds, since the deposition rate and CVs may require refinement.

We identified three categories of egress pathways (Figs. 4 and 5): a “back door” opening under the sensor loop (residues 71-77), known as the “N111-R177 backdoor” pathway(*8*, *20*, *21*) (green arrows; Fig. 4); a “back door” opening above the sensor loop known as the “R183 backdoor” pathway(*8*) (purple arrows; Fig. 4); and a “front door” opening near K18 (orange arrow; Fig. 4E). Figure 5 shows the details of each egress pathway.

**Figure 5.**
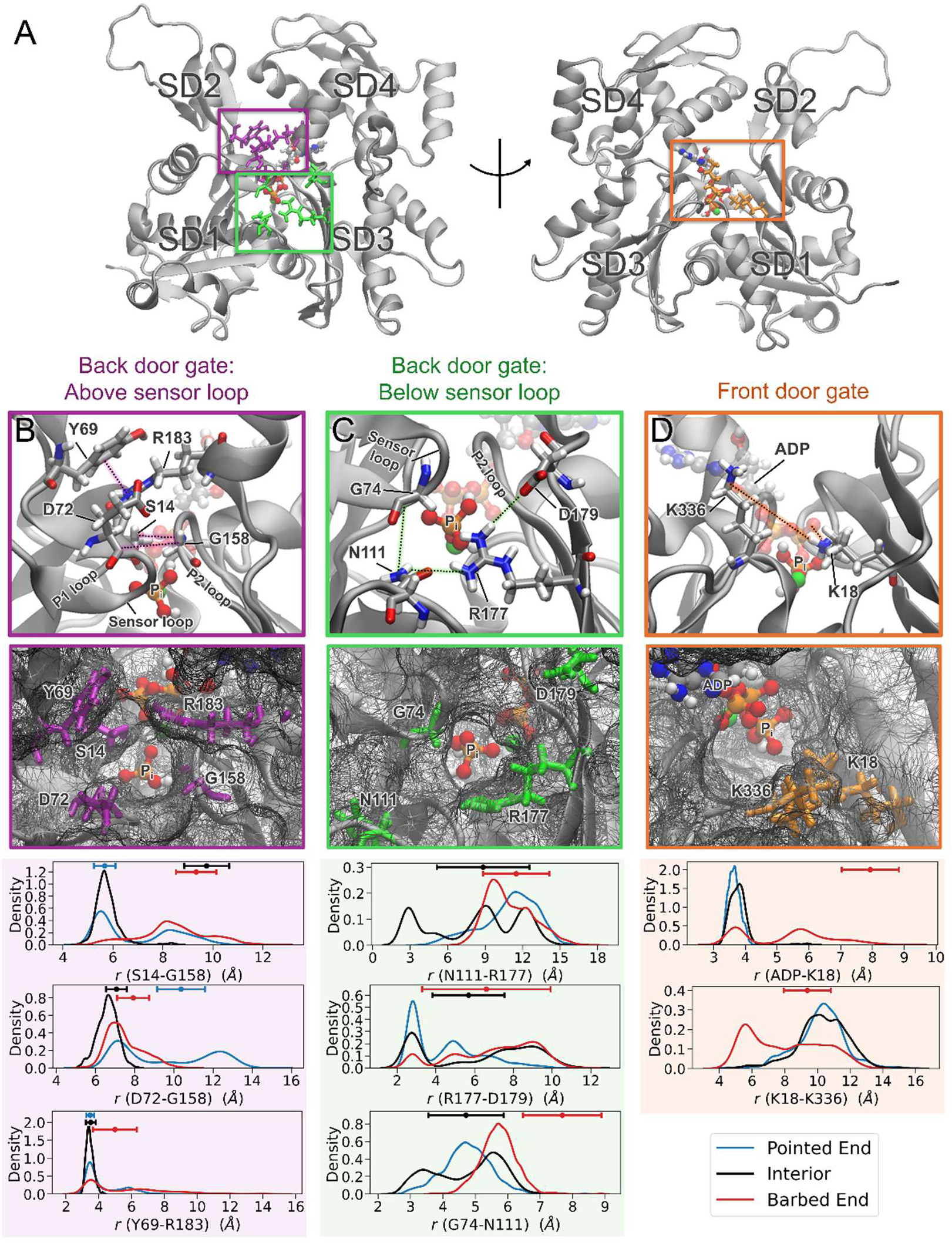
Structure and dynamics of protein residues that gate three categories of egress pathways. (A) Ribbon diagrams of the actin subunit (after 200 ns of equilibration) highlighting the 3 channels through which P_i_ egressed during MD simulations. (B-D) Columns with the three types of egress pathways. The top row has models of zoomed-in views of the amino acids that occlude the three egress pathways (licorice representation). The second row shows snapshots from outside the protein of open conformations of the channels in the barbed end subunit during unbiased simulations. The dark grey wire mech represents the protein surface with colored licorice diagrams of the relevant amino acids from the top row of panels. The graphs below plot two types of parameters for pointed end (blue), interior (black) and barbed end (red) subunits as functions of distances between the residues named on the x-axes. The smooth curves are plots of population distributions of the distances between the residues during 1.6 μs of unbiased simulations. Range bars at the top of each graph show the average (± standard deviations) of the distances at times when P_i_ used the pathway during the last 5 ns of the WT-MetaD egress simulations. (B column) Back door passage above the sensor loop (“R183 backdoor”). Distances plotted in the graphs were computed between the following atoms: S14 (CA), G158 (CA), D72 (CA), Y69 (COM of CG CD1 CD2 CE1 CE2 CZ), R183 (COM of NE, NH1, NH2). (C column) Back door passage “below” the sensor loop (“R177 backdoor”). Distances plotted in the graphs were computed between the following atoms: R177 (NE, NH1, NH2), N111 (ND2, OD1), D179 (OD1, OD2), G74 (O). (D column) Front door passage near K18. Distances plotted in the graphs were computed between the following atoms: ADP (PA, PB), K18 (NZ), K336 (NZ).

The channels are dynamic during unbiased MD simulations at 310 K, so the distances between key amino acids in each channel fluctuate on the nanosecond timescale (Fig. 5B-D). One can evaluate the plausibility of each pathway by comparing the probabilities of these distances with the range bars at the top of each graph, which show the average (and standard deviation) of the distances that permit passage of P_i_ in WT-MetaD simulations that use each egress pathway. In most but not all cases, the range bars overlap with distributions observed in unbiased simulations, indicating the channel opens on the timescales of our simulations.

### Egress from interior subunits

In 7 of 8 simulations, P_i_ exited through a pathway below the sensor loop in internal subunits (green arrow pathway, Fig. 4B; shown in more detail in Fig. 5C). This is similar to the “N111-R177 backdoor”,(*8*) although we opt for broader classes, because P_i_ can exit under the sensor loop at varying distances from R177. In 5 simulations, P_i_ first moved between A108 and H161, toward R177. In 4 of these simulations, (positively-charged) R177 interacted with (negatively-charged) P_i_, guiding its passage through A108 and H161 similar to the shuttle mechanism proposed by Wriggers and Schulten.(*20*) In the other simulation, the R177-N111 gate was wide open and R177 did not play a role in facilitating egress. In 2 simulations, P_i_ moved around A108 (toward E107) and exited under the sensor loop near residues L67 and N115 rather than through the N111-R177 opening. The N111-R177 gates of interior subunits were closed (*r* < 3.4 Å) only ∼21% of the time during 1.6 μs of simulation and favored open distances of ∼5, 9 and 12 Å (Fig. 5C upper graph) similar to that found in previous simulations.(*21*) A large range of N111-R177 distances permitted P_i_ release in WT-MetaD simulations (black bar in Fig. 5C upper panel).

In just 1 of 8 simulations, P_i_ exited above the sensor loop (purple pathway). This required S14 to separate from G158 by distances larger (range bars in Fig. 5B) than observed in the distribution of distances during 1.6 μs of unbiased simulation, making use of this pathway less likely than the “back door” pathways under the sensor loop.

### Egress from the terminal pointed end subunit

In all 8 WT-MetaD simulations, P_i_ exited above the sensor loop (purple pathway) near the “R183 backdoor”(*8*) of the terminal pointed end subunit (Figs. 4A and 5). We classify this pathway as “above sensor loop,” since the interaction between R183-Y69 need not be disrupted to open this pathway (see range bars in bottom panel of Fig. 5B). This pathway is open with either *r* (S14-G158) > 9 Å or *r* (D72-G158) > 9 Å during 62% of the 1.6 μs unbiased simulation (Fig. 5B, blue distributions). The broad range of SD2-SD1-SD3-SD4 dihedral and scissors angles sampled by the P subunit (Fig. 3B-C) allows large fluctuations in the sensor loop region (Fig. S17) and opens the exit passage by occasionally disrupting interactions between S14-G158, R183-Y69, and D72-G158 (Fig. 5B). Figs. S18-19 compares the normal modes and “above sensor loop” pathway of subunit P-1 to subunit P.

### Egress from the barbed end of the filament

We observed three plausible egress pathways from the terminal barbed end subunit (Fig. 4E): (1) In 4 of 8 simulations, P_i_ exited the subunit above the sensor loop (purple), as also observed for the terminal pointed end subunit. Along this route P_i_ interacted closely with S14 and G158 before moving out of the protein between D72, S14, and G158. Positively-charged R183 facilitated the movement through this region by interacting with P_i_ as it passed between S14 and G158, akin to the shuttling mechanism by R177 under the sensor loop. This route may be preferred in subunit B due to the slightly more twisted range of negative dihedral angles (Fig. 3B), greater fluctuations in the scissor angle of SD2 and SD4 (Fig. 3C), and, correspondingly, larger separations between S14 and G158 (Fig. 5B) than interior subunits in unbiased simulations. (2) In 3 of 8 simulations, P_i_ exited through a front door pathway (orange arrow in Fig. 4). In these cases, the interaction between K18 and ADP opens (Fig. 5D), allowing space for P_i_ to leave without unphysical distortions ADP or the protein. This pathway has not been identified previously and is only used by the barbed end subunit. The interaction of K18 with ADP also stabilizes the position of the P1 loop, which contributes to larger fluctuations in the distance between S14-G158 (Fig. 5B). (3) In 1 of 8 simulations P_i_ exited using the pathway under the sensor loop near residues I75 and I76 (green), far from the N111-R177 gate. The range bars in the bottom panel of Fig. 5C indicate this route required separations of G74 and N111 larger than observed in distributions from unbiased MD simulations, so this route is more rarely used.

We also investigated the egress pathways for the B-1 and B-2 subunits with bound ADP-P_i_ (Fig. 4C-D). P_i_ egressed from the B-1 subunit through the back door above the sensor loop in all 4 simulations, partially facilitated by R183. In 2 of these 4 simulations, it egressed between the P1 and P2 loops, slightly further from the sensor loop. Overall, the egress from the B-1 subunit is quite similar to the terminal barbed end subunit B, both with free SD1/SD3 subdomains. On the other hand, P_i_ egressed from the subunit B-2 by the back door pathway above the sensor loop in 1 of 4 simulations and under the sensor loop in 3 of 4 simulations, similar to interior subunits.

## Discussion

Our extensive unbiased and biased molecular dynamics simulations explain why the rate of P_i_ release from actin filaments is so slow and how the conditions modulate the rate by three orders of magnitude. MD simulations provide a unique perspective that complements insights from previous experiments on these systems, offering mechanistic details that are often inaccessible by structural and biochemical methods. Slow dissociation of phosphate from Mg^2+^ limits the rate of phosphate release from subunits at the ends and in the interior of filaments composed of ADP-P_i_ actin. At physiological temperature, intermittent barriers along egress channels and closed protein gates are not limiting factors though they contribute marginally to the release rate. These quantitative measurements revise previous interpretations of the P_i_ release mechanism and establish a framework for assessing how other factors, such as actin-binding proteins, may influence this important process in actin and other ATPases.

### Rate of phosphate dissociation from Mg^2+^ in water

Wriggers and Schulten(*20*) and Wang et al.(*21*) proposed that dissociation of P_i_ from the active site divalent cation is the rate-determining step in the release of P_i_ from actin subunits. These studies, however, did not investigate the faster rate of P_i_ release at the ends of the filament. Here we used extensive MD simulations with enhanced sampling methods to reevaluate this reaction under other realistic conditions including both ends of the filament.

We recognize that the complex atomic interactions governing ion pair dissociation (i.e., polarization, charge transfer, repulsion, etc.) make a quantitatively accurate estimate of the *absolute* barrier height challenging to obtain with computational methods at present. Therefore, we evaluated dissociation of P_i_ from Mg^2+^ in water to use as a point of comparison for their dissociation when bound to actin. Experimental measurements by others showed that ion pairs similar to P_i_-Mg^2+^ exist predominantly in the CIP state(*38–40*) in water and transition to the SSIP state by surmounting an activation barrier of ∼13 kcal/mol due to strong ion-ion interactions before dissociating.(*41*) To our knowledge, no chemical measurements are available for dissociation of phosphate from Mg^2+^ in water.

Our well-tempered metadynamics simulations of the CIP-to-SSIP transition of ADP-P_i_-Mg^2+^ in water yielded a barrier height of ∼21 kcal/mol and a dissociation rate constant of 0.0014 s^-1^ for P_i_ from ADP-Mg^2+^ (Fig. 2A). The height of this barrier in the simulations is likely to be too high, as expected from observations in the literature that nonpolarizable interaction potentials overstabilize CIPs.(*42–45*) However, we used the AMOEBA potential to show in the SI Section 1 and Fig. S4 that polarization effects decrease the barrier, though the absolute barrier height is sensitive to the parametrization. For these reasons, we focused on *relative* rates of CIP-to-SSIP transition in the actin systems.

### Relative rates of CIP-to-SSIP transition in actin subunits

Our calculations showed that CIP-to-SSIP transition in the barbed end subunit is 48-fold faster than in the interior subunit. This differs only 5-fold from experimental measurements by Jégou et al.,(*7*) so within one order of magnitude. The rate of P_i_ release at the pointed end has not been explicitly measured but was inferred to be much faster than the interior subunits by Fujiwara et al.(*3*) This is consistent with our calculations which indicated that the ion dissociation is 83-fold faster from the pointed end than interior subunits.

Filaments with bound jasplakinolide are an extreme example among subunits we studied. The interior subunits exhibited the slowest transition of Mg^2+^-P_i_ from CIP to SSIP, corresponding to a residence time of 1.5 – 5.0 hours. Jasplakinolide strongly inhibited P_i_ release from newly polymerized actin filaments in biochemical experiments.(*17*) After 25 minutes, only ∼5% of total P_i_ was released. Although the rate constant was not measured, jasplakinolide reduced the rate of P_i_ release by at least an order of magnitude.

Our well-tempered metadynamics simulations yielded absolute P_i_ release rates from actin that are slower (with barriers ∼20-30% too high) than biochemical experiments (Table 1). This difference stems from the empirical nature of the force field, which overstabilizes the ion pair. Further, the relative rates are slightly underestimated by about 6-fold. Electronic polarization effects may augment the relative rates of CIP to SSIP by tuning the equilibrium ion-ion distance in response to the environment more sensitively than nonpolarizable potentials. This could further increase the ion-ion interaction in environments with low effective dielectric constants and vice versa, thereby enhancing the barrier height changes in different effective dielectric environments.

Wang et al.(*21*) reported a barrier height 2-4 kcal/mol lower than ours for dissociation of Mg^2+^-P_i_ in a filament of five ADP-P_i_-actin subunits. They used the distance *r* (Mg^2+^-P_i_) as the lone CV to obtain the free energy barrier from the PMF. Our SI Section 1 shows that a second CV describing the solvent arrangement around Mg^2+^ is important to obtain the free energy barrier from the PMF, since the CIP and SSIP states overlap along the *r* (Mg^2+^-P_i_) CV (Fig. S5). That said, two CVs are necessary to adequately distinguish the reactants, transition state, and products. Using the single CV alone effectively lowers the estimated barrier of the PMF by ∼ 4 kcal/mol (Fig. S6), explaining the difference with Wang et al.(*21*). Note that 1 CV is sufficient to *promote* the CIP-to-SSIP transition to estimate first passage times. The 2^nd^ CV is not necessary because bias is not applied to the transition state or SSIP state. The rate constant obtained from fitting the ensemble of first passage times (using 1 CV) matches the estimated rate constant using transition state theory using the free energy barrier (using 2 CVs), showing that these methods are produce similar results (Section S2).

### Effect of confinement in a protein cavity on the CIP-to-SSIP transition

The transition of Mg^2+^-P_i_ from CIP to SSIP is slower inside actin subunits than in water (Fig. 1C) with rates that varied by orders of magnitude between fast transitions in the pointed end subunit and very slow transitions in interior subunits with bound jasplakinolide (Table 1). These rates scale inversely with the number of water molecules near P_i_ (Fig. 2C). This scaling behavior is because water molecules are essential for facilitating the dissociation of CIPs.(*27–30*)

Several factors likely contribute to the stabilization of ion pairs in small protein cavities. The limited numbers of water molecules available to rearrange and stabilize the transition state is likely the most important factor. In addition, the SSIP requires water molecule(s) to separate Mg^2+^ and P_i_ in the confined space. Furthermore, hydrogen bond dynamics slow in small protein cavities (Fig. S9-S10), which hinders the rearrangement of the hydrogen bond network.

Naturally, the barrier height for ion dissociation depends on the intricate balance of intermolecular interactions (in this case, ion-ion, ion-water, and ion-protein interactions). As an ion pair separates, the Coulombic interaction between the ions decreases steeply and must, in part, be balanced by stabilizing interactions with the solvent or protein. Weakly hydrated protein cavities generally have much lower dielectric screening than aqueous solution(*46*) though this depends on the shape, volume, and chemical properties of the confining medium.(*47–50*) The residence time (and correspondingly, the barrier height) is high when the effective dielectric constant of the media surrounding the ion pair is low. This effect stems from Coulomb’s Law in which the dielectric constant of the medium screens the interaction between ions. In the phosphate cavity of actin subunits, the estimated barrier height for ion dissociation (relative to that in water) is an inverse function of the volume of the cavity. This implies that the cavity volume scales with the effective dielectric environment in the phosphate cavity with contributions from the amino acids near the CIP.

Prior work on model systems consisting of mixtures of water and ethanol,(*51*, *52*) methanol,(*53*, *54*) and acetone(*55*) showed that the barrier height of the CIP-to-SSIP transition depends sensitively on the composition of the solution, the polarizability of the solution,(*56*) and the degree of confinement.(*57–59*) Our work is an example of this effect and with high biological significance.

### Which factors determine the cavity volume surrounding P_i_ in actin subunits?

Several factors collectively defined the phosphate cavities in polymerized actin subunits, including the normal mode motions and the relative positions of the four loops bordering the cavity. We confirmed this quantitatively by training a non-linear regression model to predict the phosphate cavity volume given the aforementioned parameters.

Free SD2 and SD4 of subunit P and free SD1 and SD3 of subunit B affect the equilibrium conformations of the subunits, causing them to differ in distinct ways from interior subunits. The subunits at the filament ends are less resistant to the entropically-favored fluctuations in the normal modes due to the absence of lateral and longitudinal contacts. These normal mode fluctuations are coupled to the relative positions of the loops which border the phosphate cavity. More specifically, fluctuations in the scissoring motion between SD2 and SD4 is coupled with the distances between the P1-P2 loops flanking the nucleotide phosphates. The subdomain dihedral angle between the inner and outer domains of the subunit is coupled with the relative positions of P2-sensor loop between the γ-phosphate and the protein exterior and the P2-proline rich loop.

### Protein barriers in egress channels do not limit the release of phosphate from interior actin subunits

Historically, the back door route under the sensor loop and through the “N111-R177 backdoor” has received the most attention since being identified in steered MD simulations of an actin monomer by Wriggers and Schulten(*20*). WT-MetaD simulations of a filament with 5 ADP-P_i_ subunits by Wang et al.(*21*) characterized the same pathway, because they biased P_i_ towards R177. Volume-based metadynamics simulations of an actin filament with five subunits by Oosterheert et al.(*8*) identified multiple potential egress pathways, with two “backdoor” pathways being considered most feasible: through the “N111-R177 backdoor” or the “R183 backdoor”. Analysis of a 1.1 μs simulation of a 5-mer filament showed that the hydrogen bond between R177 and N111 reversibly breaks,(*8*) in agreement with other subsequent simulations.(*21*, *25*) By inspecting the channel in a single MD frame, the former authors concluded that the channel does not fully open.

Mutation N111S partially opened the N111-R177 gate in a cryo-EM structure and increased the rate of P_i_ release. Mutations in residue R183 caused minimal changes to the cryo-EM structures and increased P_i_ release to lesser extents, reducing the authors’ interest in that pathway. Oosterheert et al.(*8*) did not calculate free energy barriers for P_i_ release from filaments. Wang et al.(*21*) did such a calculation for an interior subunit and showed that dissociation of the P_i_ from the Mg^2+^ bound ADP is the rate-limiting free energy barrier (see also the earlier discussion).

Most cryo-EM structures of actin filaments showed the sidechains of N111 and R177 hydrogen-bonded to form a tightly closed gate in the interior of the filament but partially open in the terminal barbed end subunit and open in the terminal pointed end subunit.(*8*, *24*) Cryo-EM structures showed a partially open N111-R177 gate in subunits of the N111S mutant and an open gate in subunits with coronin bound,(*8*, *23*) which both increase the rate of P_i_ release. We note that the micrographs for these cryo-EM structures were taken at temperatures of about 90 K, while the behavior of the filament at 310 K has important differences.(*25*)

Our WT-MetaD simulations identified three favorable P_i_ egress pathways that are utilized to different extents by five different subunits in the middle and ends of filaments (Fig. 4). These include back door channels above or below the sensor loop and a front door channel.

The subunits at the ends of filament ends favored alternative pathways, despite the N111-R177 gate being open in the barbed and pointed end terminal subunits. These subunits exhibited larger fluctuations about low frequency normal modes including the dihedral angle between subdomains and the scissoring angle between SD2 and SD4(*36*) which opened alternative pathways in unbiased MD simulations. These changes in conformation demonstrate that interactions (with other subunits) at both the SD1/SD3 and SD2/SD4 regions stabilize the conformations of interior subunits. Our simulations suggest that the back door pathway under the sensor loop (“N111-R177 backdoor”) is *not* the predominate mechanism for P_i_ release from either filament end. On the other hand, enhanced sampling simulations of a 5 subunit filament(*8*) showed a preference for the barbed end subunit to release phosphate through the N111-R177 backdoor. The short 1 ns constant NPT equilibration time while restraining the backbone atoms to the cryo-EM coordinates and subsequent egress simulations lasting ∼10 ns were much less than the time to fully equilibrate filaments with 7-23 subunits at 310 K.(*25*)

The egress channels are dynamic at physiological temperature of 310 K, and they opened and closed during our simulations. Hydrogen bonding networks between protein residues in these channels can impede P_i_ diffusion;(*25*) however, these barriers rearrange to allow P_i_ to pass on a much faster time scale than its dissociation from Mg^2+^.(*21*) Similarly, gates such as the hydrogen bond between N111 and R177 can block channels. The N111 and R177 gate is closed in many structures of frozen actin filaments, but is open the majority of the time in simulations at 310 K in our current and former work.(*25*) The fast diffusion of P_i_ through protein channels agrees with observations from MD simulations of myosin VI,(*60*) which showed that after dissociation of P_i_ from Mg^2+^, P_i_ diffused through protein channels and out of the protein in a few microseconds.

### Taking into account CIP-to-SSIP transition rates and egress pathways, why does P_i_ dissociate at different rates from subunits in the interior and at filament ends?

CIP-to-SSIP transition rates, egress pathways, and gates all have the potential to determine the rates that P_i_ leaves the active site and departs from actin subunits, but our simulations show that the slow CIP-to-SSIP transition rate dominates over the other factors (Fig. 1C). We find no evidence of *substantial* free energy barriers during egress from SSIP to the exterior of protein (Figs. S23-S24) that could account for the very slow rates of P_i_ release from interior subunits observed in biochemical experiments. The N111-R177 gate opens and closes during our ∼1.6 μs unbiased simulations, which is consistent with simulations of 5(*21*) and 27 Mg-ADP-P_i_-filaments.(*25*) Thus, it is unlikely that the opening the N111-R177 gate is rate-limiting. Even in the case of the filament ends, the CIP-to-SSIP transition is rate-limiting and diffusion to the protein exterior proceeds through alternative pathways, despite the N111-R177 gate being open.

### Summary of conclusions

Our data provide compelling evidence that dissociation of P_i_ from Mg^2+^ is the rate-limiting step in the release of P_i_ from subunits in the interior of filaments, at the ends, and with jasplakinolide bound. Across these systems, the rates range over three orders of magnitude due to differences in the hydration of the cavity surrounding P_i_. After dissociation from Mg^2+^, protein barriers along egress pathways have a relatively minor impact on the overall rate.

### Implications for other ATPases

The release of P_i_ is crucially important for regulating other ATPases, including myosin,(*61*) Arp2/3 complex,(*62–64*) heat shock cognate protein,(*65*) DEAD-Box Protein 5,(*66*) and AAA ATPases,(*67*, *68*) among others. While rates of phosphate release from ATPases in the ADP-Mg^2+^-P_i_ state span several orders of magnitude (fastest: > 250 s^-1^,(*69*) slowest: 0.002-0.0065 s^-1^ (*3–8*)), the estimated activation barriers range from ∼14.8 to ∼22.0 kcal/mol. The low end of this range is only slightly slower than the estimated rate for dissociation of Mg^2+^ acetyl-phosphate in aqueous solution from experiments. This places an approximate upper bound of ∼3,700 s^-1^ (residence time: 0.3 ms) on the rate of P_i_ release from ATPases.(*41*) The ∼7 kcal/mol range in barrier heights could be accounted for by confinement effects (i.e., hydration of active site) and differences in the local molecular environment of the respective proteins. Our work shows how the rate of P_i_ release (from Mg^2+^) depends on local hydration. The cation in the actin active site also has a profound effect on the P_i_ release rate. Compared with Mg^2+^-ADP-P_i_-actin filaments, P_i_ is released half as fast from Ca^2+^-ADP-P_i_-actin filaments(*9*) and unmeasurably slowly from Cr^3+^-ADP-P_i_-actin filaments.(*70*) The size and charge density of the cation (Ca^2+^ vs. Mn^2+^ vs. Mg^2+^) in the active site of myosin also affects the stability of the ion pair and the rate of P_i_ dissociation.(*71*, *72*) Our study therefore provides a framework for exploring P_i_ release (and quantifying the rate of CIP to SSIP) in other ATPases.

## Methods

### Simulation set-up

A 13-mer actin filament with each subunit in the ADP-P_i_ state was prepared from the PDB 8A2S structure.(*26*) Missing residues (1–5, 46–48) were added using MODELLER.(*73*) The resolved water molecules within 6 Å of P_i_ and Mg^2+^ were retained from the cryo-EM structure. The protein was then solvated by TIP3P water molecules allowing for at least 1.1 nm between the protein and the boundary of the box in each direction. KCl was added at a concentration of 0.1 M. An energy minimization was performed followed by a gradual heating of the system from 0 to 310K over 1 ns with 1,000 kJ/mol/nm^2^ restraints on the heavy atoms. The restraints were gradually removed by sequential 0.4 ns constant NVT simulations with 500, 250, 125, and 25 kJ/mol/nm^2^ restraints on the heavy atoms. A 1 ns NVT simulation was performed without any restraints on the atoms. A 200 ns constant NPT equilibration (without restraints) was next performed to allow the filament ends to fully relax. Zsolnay et. al showed that this equilibration time was sufficient to allow the terminal ends to relax from the interior starting conformation.(*12*) From this equilibrated structure, 2 independent constant NPT simulations were spawned and run for ∼ 800 ns each (1.6 μs total). The NPT ensemble was simulated with the Parrinello-Rahman barostat(*74*) and Bussi-Parrinello velocity rescaling thermostat(*75*) to maintain a constant pressure of 1 bar and temperature of 310 K. The leap-frog integrator was used to integrate Newton’s equations of motion with an integration step of 2 fs.

The wild type actin filament was modified to the N111S mutant using the CHARMM-GUI.(*76*, *77*) The same equilibration procedure was followed. A 500 ns unbiased NPT simulation was performed after the 200 ns equilibration. Jasplakinolide was added to the 13-mer by aligning the PDB 5OOD(*78*) structure to PDB 8A2S to place the molecule on the filament. The water molecules in the active site were retained from the PDB 8A2S structure. A harmonic restraint of 50 kJ/Å^2^ between the center-of-mass of R177(CZ, NH1, NH2) and the center-of-mass of the heterocyclic ring of jasplakinolide was added for the long NPT simulation to account for the underestimation of cation-π interactions by nonpolarizable interaction potentials.(*79*)

### Force fields

The protein was modeled using the CHARMM36m force field(*80*) and the water molecules with the TIP3P force field.(*81*) The parameters for P_i_ and jasplakinolide were obtained from CGenFF.(*82*) Coulomb and Lennard-Jones interactions were calculated in real space up to a cutoff of 12 Å. Long-range interactions were computed with an Ewald summation in reciprocal space. The simulations with the nonpolarizable interaction potentials were performed with GROMACS 2020.4(*83*) patched with PLUMED 2.7.(*84*)

### Rate calculations

The bias deposited during WT-MetaD simulations can be used to calculate the time acceleration factor for a given transition. Individual first passage times for the transition from CIP to SSIP were calculated using the method outlined by Tiwary and Parrinello.(*34*) This method uses the time integral of the bias at time *t*′, *V*_i_(*t*′), across simulation *i* to estimate the time acceleration factor *α*_i_ (Equation 4).

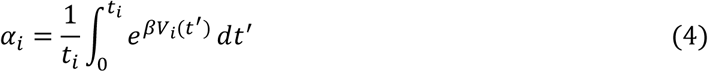

The time acceleration factors were used to rescale each biased simulation time (*t*_i_) and obtain estimates of the unbiased transition time (*α*_i_*t*_i_). Generally, 10 passage times (from independent simulations) is considered sufficient to obtain a reliable estimate of the rate. Importantly, for each independent simulation, different starting conformations were obtained at intervals of 20 ns from an unbiased simulation. This helps to account for the ensemble of conformations that exist at 310 K.(*25*) The survival probability *S*(*t*) is the fraction of independent simulations that remain in the starting CIP state after time *t*. It is assumed that *S*(*t*) = *e^-k_0_t^*, consistent with the Poisson statistics of a first-order rate. The empirical cumulative distribution function (CDF) is defined as *CDF* = 1 − *S*(*t*). The empirical CDF is fitted to obtain an estimate of *k*_O_. This estimate of the rate should be more accurate than a transition state theory estimate. Sections 1 and 2 of the SI compare the rate constants from fitting rescaled first passage times with estimates from transition state theory, showing that this method is in quantitative agreement with the free energy barrier from the PMF with 2 CVs (ion-ion distance and coordination number of Mg^2+^).

Generally, a very slow pace of bias deposition is required to ensure bias is not applied to the transition state and that the deposited bias does not exceed the barrier height. As suggested by others in the literature,(*85*) we adapt the pace at which the bias is deposited once the simulation reaches certain points on the free energy surface. This helps to decrease the computational cost by adding bias quickly when the system is far from the transition state and gradually slowing the pace once the system is close to the transition state. Specifically, the bias was deposited every 1 ps until *r* (Mg^2+^-P_i_) reached 4.0 Å. The frequency was lowered 10-fold so that the bias was deposited every 10 ps until *r* (Mg^2+^-P_i_) reached 4.15 Å. Finally, the biased was deposited every 100 ps until the SSIP state was reached (defined as *r* (Mg^2+^-P_i_) = 5.0 Å). Note that once the transition state is surmounted, the system quickly relaxes to the SSIP well without depositing bias to the SSIP state. The starting Gaussian bias height was 1 kJ/mol and width was 0.05 Å. A bias factor of 10 (14 for jasplakinolide) was used to temper the hill height over the simulation. The uncertainty in the rate was quantified using bootstrapping. Specifically, the standard error of the rate constant was obtained as the standard deviation of the log-normal distribution of rate constants from 10,000 bootstrap samples.

### Egress simulations

Simulations starting from the CIP were performed by biasing the distance between Mg^2+^ and the center-of-mass of P_i_. The pace was set to deposit bias every 10 ps. The bias height was set at 1.0 kJ/mol, width at 0.05 Å, and bias factor to 10. Importantly, these simulations do not favor a particular path by biasing the Mg^2+^-P_i_ distance alone. The addition of bias is not aggressive which results in physically plausible pathways, without noticeable distortions to the protein or ADP. Eight independent simulations were performed for each subunit (four for B-1, B-2 subunits), starting from different structures sampled every 20 ns from an unbiased NPT simulation.

### Analysis

The phosphate cavity volumes were measured using POVME3.(*35*) The center of the seed sphere was defined as the position of the phosphorus atom of P_i_. The radius of the sphere was set to 5 Å. POVME removes the volume within the seed sphere that contains protein, ADP, or Mg^2+^. Note that the phosphate cavity volumes include the volume of P_i_ and, thus, reflects the volume that P_i_ and water can exist. The subdomains of the actin subunit are defined as follows: SD1 (residues 1-32, 70-144, 338 to 375), SD2 (residues 33-69), SD3 (residues 145-180, 270-337), and SD4 (residues 181-269). A Random Forest Regressor(*37*) (implemented in Scikit-learn(*86*)) was used to train a model to predict the phosphate cavity volume given the distances between the four loops which border the cavity and the two major normal modes.

## Acknowledgements.

Research reported in this publication was supported by the National Institute of General Medical Sciences (NIGMS) of the National Institutes of Health (NIH) under award numbers F32GM161057 to K.M.H, and R35GM158238 to G.A.V. The content is solely the responsibility of the authors and does not necessarily represent the official views of the NIH. Computational resources were provided by the University of Chicago Research Computing Center and the NIH-funded Beagle-3 computer (NIH award 1S10OD028655-01).

## Author Contributions

Conceptualization: KMH, TDP, GAV

Methodology: KMH, SSI, YW

Investigation: KMH

Visualization: KMH

Supervision: TDP, GAV

Writing—original draft: KMH

Writing—review & editing: KMH, SSI, YW, TDP, GAV; all authors discussed results and contributed to manuscript revisions.

## Data, Materials, and Software Availability

Scripts and files for rate calculations, free energy calculations, and parameters measured over MD simulations are available online (https://doi.org/10.5281/zenodo.20753325). Due to the large file sizes and number of files, trajectories will be made available upon request.

## References

1. A. Wegner, Head to tail polymerization of actin. Journal of Molecular Biology 108, 139–150 (1976).

2. E. D. Korn, M.-F. Carlier, D. Pantaloni, Actin Polymerization and ATP Hydrolysis. Science 238, 638–644 (1987).

3. I. Fujiwara, D. Vavylonis, T. D. Pollard, Polymerization kinetics of ADP- and ADP-Pi-actin determined by fluorescence microscopy. Proceedings of the National Academy of Sciences 104, 8827–8832 (2007).

4. L. Blanchoin, T. D. Pollard, Mechanism of Interaction of *Acanthamoeba* Actophorin (ADF/Cofilin) with Actin Filaments*. Journal of Biological Chemistry 274, 15538–15546 (1999).

5. M.-F. Carlier, Measurement of Pi dissociation from actin filaments following ATP hydrolysis using a linked enzyme assay. Biochemical and Biophysical Research Communications 143, 1069–1075 (1987).

6. R. Melki, S. Fievez, M.-F. Carlier, Continuous Monitoring of Pi Release Following Nucleotide Hydrolysis in Actin or Tubulin Assembly Using 2-Amino-6-mercapto-7-methylpurine Ribonucleoside and Purine-Nucleoside Phosphorylase as an Enzyme-Linked Assay. Biochemistry 35, 12038–12045 (1996).

7. A. Jégou, T. Niedermayer, J. Orbán, D. Didry, R. Lipowsky, M.-F. Carlier, G. Romet-Lemonne, Individual Actin Filaments in a Microfluidic Flow Reveal the Mechanism of ATP Hydrolysis and Give Insight Into the Properties of Profilin. PLOS Biology 9, e1001161 (2011).

8. W. Oosterheert, F. E. C. Blanc, A. Roy, A. Belyy, M. B. Sanders, O. Hofnagel, G. Hummer, P. Bieling, S. Raunser, Molecular mechanisms of inorganic-phosphate release from the core and barbed end of actin filaments. Nat Struct Mol Biol 30, 1774–1785 (2023).

9. M. F. Carlier, D. Pantaloni, Direct evidence for ADP-inorganic phosphate-F-actin as the major intermediate in ATP-actin polymerization. Rate of dissociation of inorganic phosphate from actin filaments. Biochemistry 25, 7789–7792 (1986).

10. T. D. Pollard, I. Goldberg, W. H. Schwarz, Nucleotide exchange, structure, and mechanical properties of filaments assembled from ATP-actin and ADP-actin. Journal of Biological Chemistry 267, 20339–20345 (1992).

11. T. D. Pollard, L. Blanchoin, R. D. Mullins, Molecular Mechanisms Controlling Actin Filament Dynamics in Nonmuscle Cells. Annual Review of Biophysics 29, 545–576 (2000).

12. V. Zsolnay, H. H. Katkar, S. Z. Chou, T. D. Pollard, G. A. Voth, Structural basis for polarized elongation of actin filaments. Proc. Natl. Acad. Sci. U.S.A. 117, 30458–30464 (2020).

13. H. Isambert, P. Venier, A. C. Maggs, A. Fattoum, R. Kassab, D. Pantaloni, M. F. Carlier, Flexibility of actin filaments derived from thermal fluctuations. Effect of bound nucleotide, phalloidin, and muscle regulatory proteins. J Biol Chem 270, 11437–11444 (1995).

14. M. J. Reynolds, C. Hachicho, A. G. Carl, R. Gong, G. M. Alushin, Bending forces and nucleotide state jointly regulate F-actin structure. Nature 611, 380–386 (2022).

15. C. Suarez, J. Roland, R. Boujemaa-Paterski, H. Kang, B. R. McCullough, A.-C. Reymann, C. Guérin, J.-L. Martiel, E. M. De La Cruz, L. Blanchoin, Cofilin Tunes the Nucleotide State of Actin Filaments and Severs at Bare and Decorated Segment Boundaries. Current Biology 21, 862–868 (2011).

16. A. Jégou, T. Niedermayer, J. Orbán, D. Didry, R. Lipowsky, M.-F. Carlier, G. Romet-Lemonne, Individual Actin Filaments in a Microfluidic Flow Reveal the Mechanism of ATP Hydrolysis and Give Insight Into the Properties of Profilin. PLOS Biology 9, e1001161 (2011).

17. A. Vig, R. Ohmacht, É. Jámbor, B. Bugyi, M. Nyitrai, G. Hild, The effect of toxins on inorganic phosphate release during actin polymerization. Eur Biophys J 40, 619–626 (2011).

18. B. Ilkovski, S. T. Cooper, K. Nowak, M. M. Ryan, N. Yang, C. Schnell, H. J. Durling, L. G. Roddick, I. Wilkinson, A. J. Kornberg, K. J. Collins, G. Wallace, P. Gunning, E. C. Hardeman, N. G. Laing, K. N. North, Nemaline Myopathy Caused by Mutations in the Muscle α-Skeletal-Actin Gene. The American Journal of Human Genetics 68, 1333–1343 (2001).

19. S. Ladha, S. Coons, S. Johnsen, N. Sambuughin, R. Bien-Wilner, K. Sivakumar, Histopathologic Progression and a Novel Mutation in a Child With Nemaline Myopathy. J Child Neurol 23, 813–817 (2008).

20. W. Wriggers, K. Schulten, Investigating a back door mechanism of actin phosphate release by steered molecular dynamics. *Proteins: Structure*, Function, and Bioinformatics 35, 262–273 (1999).

21. Y. Wang, J. Wu, V. Zsolnay, T. D. Pollard, G. A. Voth, Mechanism of phosphate release from actin filaments. Proceedings of the National Academy of Sciences 121, e2408156121 (2024).

22. S. Z. Chou, T. D. Pollard, Mechanism of actin polymerization revealed by cryo-EM structures of actin filaments with three different bound nucleotides. Proceedings of the National Academy of Sciences 116, 4265–4274 (2019).

23. W. Oosterheert, M. B. Sanders, O. Hofnagel, P. Bieling, S. Raunser, Choreography of rapid actin filament disassembly by coronin, cofilin, and AIP1. Cell 0 (2025).

24. S. Z. Chou, T. D. Pollard, Cryo-EM structures of both ends of the actin filament explain why the barbed end elongates faster than the pointed end. bioRxiv [Preprint] (2023). 10.1101/2023.05.12.540494.

25. S. Sridharan Iyer, K. M. Herman, T. Paul, Y. Wang, T. D. Pollard, G. A. Voth, Reaching the full potential of cryo-EM reconstructions with molecular dynamics simulations at 310 K: Actin filaments as an example. Proceedings of the National Academy of Sciences 122, e2521421122 (2025).

26. W. Oosterheert, B. U. Klink, A. Belyy, S. Pospich, S. Raunser, Structural basis of actin filament assembly and aging. Nature 611, 374–379 (2022).

27. P. L. Geissler, C. Dellago, D. Chandler, Kinetic Pathways of Ion Pair Dissociation in Water. J. Phys. Chem. B 103, 3706–3710 (1999).

28. A. J. Ballard, C. Dellago, Toward the Mechanism of Ionic Dissociation in Water. J. Phys. Chem. B 116, 13490–13497 (2012).

29. R. G. Mullen, J.-E. Shea, B. Peters, Transmission Coefficients, Committors, and Solvent Coordinates in Ion-Pair Dissociation. J. Chem. Theory Comput. 10, 659–667 (2014).

30. S. Roy, M. D. Baer, C. J. Mundy, G. K. Schenter, Marcus Theory of Ion-Pairing. J. Chem. Theory Comput. 13, 3470–3477 (2017).

31. A. Barducci, G. Bussi, M. Parrinello, Well-Tempered Metadynamics: A Smoothly Converging and Tunable Free-Energy Method. Phys. Rev. Lett. 100, 020603 (2008).

32. J. F. Dama, M. Parrinello, G. A. Voth, Well-Tempered Metadynamics Converges Asymptotically. Phys. Rev. Lett. 112, 240602 (2014).

33. W. Humphrey, A. Dalke, K. Schulten, VMD: Visual molecular dynamics. Journal of Molecular Graphics 14, 33–38 (1996).

34. P. Tiwary, M. Parrinello, From Metadynamics to Dynamics. Phys. Rev. Lett. 111, 230602 (2013).

35. J. R. Wagner, J. Sørensen, N. Hensley, C. Wong, C. Zhu, T. Perison, R. E. Amaro, POVME 3.0: Software for Mapping Binding Pocket Flexibility. J. Chem. Theory Comput. 13, 4584–4592 (2017).

36. M. M. Tirion, D. ben-Avraham, Normal Mode Analysis of G-actin. Journal of Molecular Biology 230, 186–195 (1993).

37. L. Breiman, Random Forests. Machine Learning 45, 5–32 (2001).

38. J. Schauss, F. Dahms, B. P. Fingerhut, T. Elsaesser, Phosphate–Magnesium Ion Interactions in Water Probed by Ultrafast Two-Dimensional Infrared Spectroscopy. J. Phys. Chem. Lett. 10, 238–243 (2019).

39. J. Schauss, A. Kundu, B. P. Fingerhut, T. Elsaesser, Contact Ion Pairs of Phosphate Groups in Water: Two-Dimensional Infrared Spectroscopy of Dimethyl Phosphate and ab Initio Simulations. J. Phys. Chem. Lett. 10, 6281–6286 (2019).

40. B. Kutus, K. Wagner, M. Wagner, J. Hunger, Ion-pairing equilibria and kinetics of dimethyl phosphate: A model for counter-ion binding to the phosphate backbone of nucleic acids. Journal of Molecular Liquids 363, 119868 (2022).

41. J. A. Cowan, Metallobiochemistry of magnesium. Coordination complexes with biological substrates: site specificity, kinetics and thermodynamics of binding, and implications for activity. Inorg. Chem. 30, 2740–2747 (1991).

42. S. Falkner, N. Schwierz, Kinetic pathways of water exchange in the first hydration shell of magnesium: Influence of water model and ionic force field. J. Chem. Phys. 155, 084503 (2021).

43. E. Wernersson, P. Jungwirth, Effect of Water Polarizability on the Properties of Solutions of Polyvalent Ions: Simulations of Aqueous Sodium Sulfate with Different Force Fields. J. Chem. Theory Comput. 6, 3233–3240 (2010).

44. E. Pluhařová, P. E. Mason, P. Jungwirth, Ion Pairing in Aqueous Lithium Salt Solutions with Monovalent and Divalent Counter-Anions. J. Phys. Chem. A 117, 11766–11773 (2013).

45. E. Wernersson, P. Jungwirth, Effect of Water Polarizability on the Properties of Solutions of Polyvalent Ions: Simulations of Aqueous Sodium Sulfate with Different Force Fields. J. Chem. Theory Comput. 6, 3233–3240 (2010).

46. T. Simonson, C. L. Brooks, Charge Screening and the Dielectric Constant of Proteins: Insights from Molecular Dynamics. J. Am. Chem. Soc. 118, 8452–8458 (1996).

47. E. Papadopoulou, J. Zavadlav, R. Podgornik, M. Praprotnik, P. Koumoutsakos, Tuning the Dielectric Response of Water in Nanoconfinement through Surface Wettability. ACS Nano, doi: 10.1021/acsnano.1c08512 (2021).

48. S. Senapati, A. Chandra, Dielectric Constant of Water Confined in a Nanocavity. J. Phys. Chem. B 105, 5106–5109 (2001).

49. I. Ahmadabadi, A. Esfandiar, A. Hassanali, M. R. Ejtehadi, Structural and dynamical fingerprints of the anomalous dielectric properties of water under confinement. Phys. Rev. Mater. 5, 024008 (2021).

50. S. Mondal, S. Acharya, B. Bagchi, Altered polar character of nanoconfined liquid water. Phys. Rev. Res. 1, 033145 (2019).

51. A. Chatterjee, M. K. Dixit, B. L. Tembe, Solvation Structures and Dynamics of the Magnesium Chloride (Mg2+–Cl–) Ion Pair in Water–Ethanol Mixtures. J. Phys. Chem. A 117, 8703–8709 (2013).

52. M. D. Meti, M. Dixit, T. Hajari, B. L. Tembe, Ion pairing and preferential solvation of butylmethylimidazolium chloride ion pair in water-ethanol mixtures by using molecular dynamics simulations. Chemical Physics Letters 720, 107–112 (2019).

53. M. Kelley, A. Donley, S. Clark, A. Clark, Structure and Dynamics of NaCl Ion Pairing in Solutions of Water and Methanol. J. Phys. Chem. B 119, 15652–15661 (2015).

54. S. Keshri, A. Sarkar, B. L. Tembe, Molecular Dynamics Simulation of Na+–Cl– Ion-Pair in Water–Methanol Mixtures under Supercritical and Ambient Conditions. J. Phys. Chem. B 119, 15471–15484 (2015).

55. A. A. Siddique, M. K. Dixit, B. L. Tembe, Molecular dynamics simulations of Ca2+Cl− ion pair in polar mixtures of acetone and water: Solvation and dynamical studies. Chemical Physics Letters 662, 306–316 (2016).

56. H. Gudla, Y. Shao, S. Phunnarungsi, D. Brandell, C. Zhang, Importance of the Ion-Pair Lifetime in Polymer Electrolytes. J. Phys. Chem. Lett. 12, 8460–8464 (2021).

57. K. D. Fong, B. Sumić, N. O’Neill, C. Schran, C. P. Grey, A. Michaelides, The Interplay of Solvation and Polarization Effects on Ion Pairing in Nanoconfined Electrolytes. Nano Lett. 24, 5024–5030 (2024).

58. R. Wang, P. Tiwary, Atomic scale insights into NaCl nucleation in nanoconfined environments. doi: 10.1039/D4SC04042B (2024).

59. B. T. Fashina, A. P. Baldo, H. Watts, K. Leung, J. D. Kubicki, A. G. Ilgen, Enhanced ion pairing of calcium, zinc, and lanthanum with acetate in silica nanopores. Microporous and Mesoporous Materials 387, 113538 (2025).

60. M. L. Mugnai, D. Thirumalai, Step-Wise Hydration of Magnesium by Four Water Molecules Precedes Phosphate Release in a Myosin Motor. J. Phys. Chem. B 125, 1107–1117 (2021).

61. Y. Takagi, H. Shuman, Y. E. Goldman, Coupling between phosphate release and force generation in muscle actomyosin. Philos Trans R Soc Lond B Biol Sci 359, 1913–1920 (2004).

62. N. G. Pandit, W. Cao, J. Bibeau, E. M. Johnson-Chavarria, E. W. Taylor, T. D. Pollard, E. M. De La Cruz, Force and phosphate release from Arp2/3 complex promote dissociation of actin filament branches. Proceedings of the National Academy of Sciences 117, 13519–13528 (2020).

63. S. S. Chavali, S. Z. Chou, W. Cao, T. D. Pollard, E. M. De La Cruz, C. V. Sindelar, Cryo-EM structures reveal how phosphate release from Arp3 weakens actin filament branches formed by Arp2/3 complex. Nat Commun 15, 2059 (2024).

64. J. Xiao, R. Pagès, G. Romet-Lemonne, A. Jégou, Inorganic phosphate in Arp2/3 complex acts as a rapid switch for the stability of actin filament branches. bioRxiv [Preprint] (2025). 10.1101/2025.07.18.665541.

65. J. H. Ha, D. B. McKay, ATPase kinetics of recombinant bovine 70 kDa heat shock cognate protein and its amino-terminal ATPase domain. Biochemistry 33, 14625–14635 (1994).

66. E. V. Wong, W. Cao, J. Vörös, M. Merchant, Y. Modis, D. D. Hackney, B. Montpetit, E. M. De La Cruz, Pi Release Limits the Intrinsic and RNA-Stimulated ATPase Cycles of DEAD-Box Protein 5 (Dbp5). Journal of Molecular Biology 428, 492–508 (2016).

67. I. Rouiller, B. DeLaBarre, A. P. May, W. I. Weis, A. T. Brunger, R. A. Milligan, E. M. Wilson-Kubalek, Conformational changes of the multifunction p97 AAA ATPase during its ATPase cycle. Nat Struct Mol Biol 9, 950–957 (2002).

68. X. Liu, L. Rao, A. Gennerich, The regulatory function of the AAA4 ATPase domain of cytoplasmic dynein. Nat Commun 11, 5952 (2020).

69. E. M. De La Cruz, A. L. Wells, S. S. Rosenfeld, E. M. Ostap, H. L. Sweeney, The kinetic mechanism of myosin V. Proceedings of the National Academy of Sciences 96, 13726–13731 (1999).

70. C. Valentin-Ranc, M. F. Carlier, Evidence for the Direct Interaction Between Tightly Bound Divalent Metal Ion and ATP on Actin. Journal of Biological Chemistry 264, 20871–20880 (1989).

71. Y. V. Tkachev, J. Ge, I. V. Negrashov, Y. E. Nesmelov, Metal cation controls myosin and actomyosin kinetics. Protein Science 22, 1766–1774 (2013).

72. J. Ge, F. Huang, Y. E. Nesmelov, Metal cation controls phosphate release in the myosin ATPase. Protein Science 26, 2181–2186 (2017).

73. B. Webb, A. Sali, Comparative Protein Structure Modeling Using MODELLER. Current Protocols in Bioinformatics 54, 5.6.1–5.6.37 (2016).

74. M. Parrinello, A. Rahman, Polymorphic transitions in single crystals: A new molecular dynamics method. J. Appl. Phys. 52, 7182–7190 (1981).

75. G. Bussi, D. Donadio, M. Parrinello, Canonical sampling through velocity rescaling. J. Chem. Phys. 126, 014101 (2007).

76. S. Jo, T. Kim, V. G. Iyer, W. Im, CHARMM-GUI: A web-based graphical user interface for CHARMM. Journal of Computational Chemistry 29, 1859–1865 (2008).

77. S. Jo, X. Cheng, S. M. Islam, L. Huang, H. Rui, A. Zhu, H. S. Lee, Y. Qi, W. Han, K. Vanommeslaeghe, A. D. MacKerell, B. Roux, W. Im, “Chapter Eight - CHARMM-GUI PDB Manipulator for Advanced Modeling and Simulations of Proteins Containing Nonstandard Residues” in Advances in Protein Chemistry and Structural Biology, T. Karabencheva-Christova, Ed. (Academic Press, 2014; https://www.sciencedirect.com/science/article/pii/S1876162314000030)vol. 96 of *Biomolecular Modelling and Simulations*, pp. 235–265.

78. F. Merino, S. Pospich, J. Funk, T. Wagner, F. Küllmer, H.-D. Arndt, P. Bieling, S. Raunser, Structural transitions of F-actin upon ATP hydrolysis at near-atomic resolution revealed by cryo-EM. Nat Struct Mol Biol 25, 528–537 (2018).

79. H. Liu, H. Fu, X. Shao, W. Cai, C. Chipot, Accurate Description of Cation−π Interactions in Proteins with a Nonpolarizable Force Field at No Additional Cost. J. Chem. Theory Comput. 16, 6397–6407 (2020).

80. J. Huang, S. Rauscher, G. Nawrocki, T. Ran, M. Feig, B. L. de Groot, H. Grubmüller, A. D. MacKerell, CHARMM36m: an improved force field for folded and intrinsically disordered proteins. Nat Methods 14, 71–73 (2017).

81. A. D. Jr. MacKerell, D. Bashford, M. Bellott, R. L. Jr. Dunbrack, J. D. Evanseck, M. J. Field, S. Fischer, J. Gao, H. Guo, S. Ha, D. Joseph-McCarthy, L. Kuchnir, K. Kuczera, F. T. K. Lau, C. Mattos, S. Michnick, T. Ngo, D. T. Nguyen, B. Prodhom, W. E. Reiher, B. Roux, M. Schlenkrich, J.C. Smith, R. Stote, J. Straub, M. Watanabe, J. Wiórkiewicz-Kuczera, D. Yin, M. Karplus, All-Atom Empirical Potential for Molecular Modeling and Dynamics Studies of Proteins. J. Phys. Chem. B 102, 3586–3616 (1998).

82. K. Vanommeslaeghe, E. Hatcher, C. Acharya, S. Kundu, S. Zhong, J. Shim, E. Darian, O. Guvench, P. Lopes, I. Vorobyov, A. D. Mackerell Jr., CHARMM general force field: A force field for drug-like molecules compatible with the CHARMM all-atom additive biological force fields. Journal of Computational Chemistry 31, 671–690 (2010).

83. M. J. Abraham, T. Murtola, R. Schulz, S. Páll, J. C. Smith, B. Hess, E. Lindahl, GROMACS: High performance molecular simulations through multi-level parallelism from laptops to supercomputers. SoftwareX 1–2, 19–25 (2015).

84. M. Bonomi, D. Branduardi, G. Bussi, C. Camilloni, D. Provasi, P. Raiteri, D. Donadio, F. Marinelli, F. Pietrucci, R. A. Broglia, M. Parrinello, PLUMED: A portable plugin for free-energy calculations with molecular dynamics. Computer Physics Communications 180, 1961–1972 (2009).

85. Y. Wang, O. Valsson, P. Tiwary, M. Parrinello, K. Lindorff-Larsen, Frequency adaptive metadynamics for the calculation of rare-event kinetics. J. Chem. Phys. 149, 072309 (2018).

86. F. Pedregosa, G. Varoquaux, A. Gramfort, V. Michel, B. Thirion, O. Grisel, M. Blondel, P. Prettenhofer, R. Weiss, V. Dubourg, Scikit-learn: Machine learning in Python. the Journal of machine Learning research 12, 2825–2830 (2011).

87. K. M. Herman, A. J. Stone, S. S. Xantheas, A classical model for three-body interactions in aqueous ionic systems. J. Chem. Phys. 157, 024101 (2022).

88. K. M. Herman, A. J. Stone, S. S. Xantheas, Accurate Calculation of Many-Body Energies in Water Clusters Using a Classical Geometry-Dependent Induction Model. J. Chem. Theory Comput. 19, 6805–6815 (2023).

89. C. Zhang, F. Giberti, E. Sevgen, J. J. de Pablo, F. Gygi, G. Galli, Dissociation of salts in water under pressure. Nat Commun 11, 3037 (2020).

90. J. Zhang, L. Bonati, E. Trizio, O. Zhang, Y. Kang, T. Hou, M. Parrinello, Descriptor-Free Collective Variables from Geometric Graph Neural Networks. J. Chem. Theory Comput. 20, 10787–10797 (2024).

91. R. Caminiti, G. Licheri, G. Piccaluga, G. Pinna, X-ray diffraction study of a “three-ion” aqueous solution. Chemical Physics Letters 47, 275–278 (1977).

92. N. A. Matwiyoff, H. Taube, Direct determination of the solvation number of magnesium(II) ion in water, aqueous acetone, and methanolic acetone solutions. J. Am. Chem. Soc. 90, 2796–2800 (1968).

93. C. C. Pye, W. W. Rudolph, An ab Initio and Raman Investigation of Magnesium(II) Hydration. J. Phys. Chem. A 102, 9933–9943 (1998).

94. W. E, W. Ren, E. Vanden-Eijnden, Simplified and improved string method for computing the minimum energy paths in barrier-crossing events. J. Chem. Phys. 126, 164103 (2007).

95. A. Bleuzen, P.-A. Pittet, L. Helm, A. E. Merbach, Water exchange on magnesium(II) in aqueous solution: a variable temperature and pressure 17O NMR study. Magnetic Resonance in Chemistry 35, 765–773 (1997).

96. J. A. Cowan, H.-W. Huang, L.-Y. Hsu, Sequence selective coordination of Mg2+(aq) to DNA. Journal of Inorganic Biochemistry 52, 121–129 (1993).

97. J. A. Cowan, Coordination chemistry of magnesium ions and 5S rRNA (Escherichia coli): binding parameters, ligand symmetry, and implications for activity. J. Am. Chem. Soc. 113, 675–676 (1991).

98. O. Allnér, L. Nilsson, A. Villa, Magnesium Ion–Water Coordination and Exchange in Biomolecular Simulations. J. Chem. Theory Comput. 8, 1493–1502 (2012).

99. Y. Shi, Z. Xia, J. Zhang, R. Best, C. Wu, J. W. Ponder, P. Ren, Polarizable Atomic Multipole-Based AMOEBA Force Field for Proteins. J. Chem. Theory Comput. 9, 4046–4063 (2013).

100. C. Zhang, C. Lu, Z. Jing, C. Wu, J.-P. Piquemal, J. W. Ponder, P. Ren, AMOEBA Polarizable Atomic Multipole Force Field for Nucleic Acids. J. Chem. Theory Comput. 14, 2084–2108 (2018).

101. B. Walker, Z. Jing, P. Ren, Molecular dynamics free energy simulations of ATP:Mg2+ and ADP:Mg2+ using the polarisable force field AMOEBA. Molecular Simulation 47, 439–448 (2021).

102. M. Kumar, T. Simonson, G. Ohanessian, C. Clavaguéra, Structure and Thermodynamics of Mg:Phosphate Interactions in Water: A Simulation Study. ChemPhysChem 16, 658–665 (2015).

103. J. M. Delgado, P. R. Nagy, S. Varma, Polarizable AMOEBA Model for Simulating Mg2+·Protein·Nucleotide Complexes. J Chem Inf Model 64, 378–392 (2024).

104. O. Adjoua, L. Lagardère, L.-H. Jolly, A. Durocher, T. Very, I. Dupays, Z. Wang, T. J. Inizan, F. Célerse, P. Ren, J. W. Ponder, J.-P. Piquemal, Tinker-HP: Accelerating Molecular Dynamics Simulations of Large Complex Systems with Advanced Point Dipole Polarizable Force Fields Using GPUs and Multi-GPU Systems. J. Chem. Theory Comput. 17, 2034–2053 (2021).

105. L. Lagardère, L.-H. Jolly, F. Lipparini, F. Aviat, B. Stamm, Z. F. Jing, M. Harger, H. Torabifard, G. Andrés Cisneros, M. J. Schnieders, N. Gresh, Y. Maday, P. Y. Ren, J. W. Ponder, J.-P. Piquemal, Tinker-HP: a massively parallel molecular dynamics package for multiscale simulations of large complex systems with advanced point dipole polarizable force fields. Chemical Science 9, 956–972 (2018).

106. P. Tiwary, M. Parrinello, A Time-Independent Free Energy Estimator for Metadynamics. J. Phys. Chem. B 119, 736–742 (2015).

107. H. Eyring, The Activated Complex in Chemical Reactions. J. Chem. Phys. 3, 107–115 (1935).

108. M. G. Evans, M. Polanyi, Some applications of the transition state method to the calculation of reaction velocities, especially in solution. Transactions of the Faraday Society 31, 875–894 (1935).

109. G. K. Schenter, B. C. Garrett, D. G. Truhlar, Generalized transition state theory in terms of the potential of mean force. The Journal of chemical physics 119, 5828 (2003).

110. Y. Liu, C. Li, G. A. Voth, Generalized Transition State Theory Treatment of Water-Assisted Proton Transport Processes in Proteins. J. Phys. Chem. B 126, 10452–10459 (2022).

