## Supplemental Information for "Physical Confinement Modulates the Rate-Limiting Transition in the Release of Phosphate from Actin Filaments"

### Table of Contents

**Section 1.** Thermodynamics of the transition of the contact ion pair (CIP) to solvent-separated ion pair (SSIP) for ADP-$Mg^{2+}$-$P_i$ in water

**Section 2.** Kinetics of the transition of CIP to SSIP for ADP-$Mg^{2+}$-$P_i$ in water

**Section 3.** Ion pair dissociation in the phosphate cavity of actin filaments

**Section 4.** Hydrogen bond lifetime in the protein cavity near the CIP

**Section 5.** Relationship between protein conformations and phosphate cavity volumes

**Section 6.** Comparison of conformation of subunit P-1 to subunit P

**Section 7.** Effects of mutations on $P_i$ release

**Section 8.** Thermodynamics of $P_i$ release pathways from different subunits

### Section 1: Thermodynamics of the transition of the contact ion pair (CIP) to solvent-separated ion pair (SSIP) for ADP-$Mg^{2+}$-$P_i$ in water

Past work established the complexities of ion pairing including the importance of solvent molecules in facilitating the transition and the role of many-body effects arising from polarization.(*27–30*, *57*, *87*, *88*) The below work on the ion pair (Fig. S1) serves as a benchmark for the computational methods, provides insights into the mechanism of ion pair dissociation, and is a point of comparison for the rates of ion pair dissociation in actin filaments.

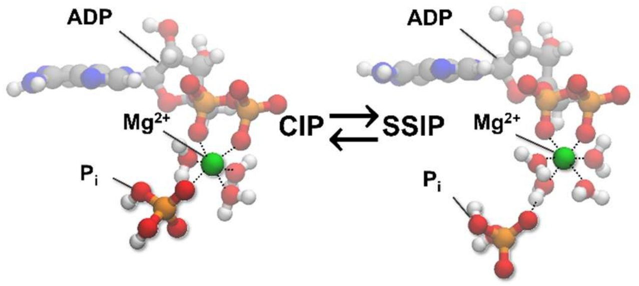


**Figure S1.** CPK models of the transition from the contact ion pair (CIP) with a phosphate oxygen bound directly to $Mg^{2+}$ to the solvent-separated ion pair (SSIP) states of ADP-$Mg^{2+}$-$P_i$ which was proposed to be the rate-determining step for phosphate release in previous works.(*20*, *21*)

The CIP (ADP-$Mg^{2+}$-$P_i$) was equilibrated in a TIP3P water box with at least 1.1 nm separating the CIP from the edge of the box in all directions. A 5 ns NPT simulation was performed to equilibrate the CIP.

We used well-tempered metadynamics (WT-MetaD), an enhanced sampling method, to simulate the dissociation of $P_i$ from ADP-$Mg^{2+}$ in a box of 1,216 water molecules. WT-MetaD(*31*, *32*) accelerates sampling along select degrees of freedom, termed collective variables (CVs), by depositing Gaussian biases along these CVs throughout an MD simulation. After sampling the transition from CIP to SSIP (and back) several times in a WT-MetaD simulation, we used the biases deposited over the simulation to reweight the probability density (of the CV) to obtain a potential of mean force (PMF).

We calculated the 2-dimensional potential of mean force (PMF) for the transition from CIP (ADP-$Mg^{2+}$-$P_i$) to SSIP (ADP-$Mg^{2+}\cdots P_i$) in water (Fig. S2A). The salient degrees of freedom for this transition are (x-axis) the distance between the centers-of-mass of $Mg^{2+}$ and $P_i$, $r$ ($P_i$-$Mg^{2+}$), and (y-axis) the number of oxygen atoms of $H_2O$ and ADP coordinated with $Mg^{2+}$, $CN$ ($Mg^{2+}$). The latter is defined as the coordination number of $Mg^{2+}$ with oxygen atoms of ADP and $H_2O$ (S1):

$$CN\ (Mg^{2+}) = \sum_{\substack{Oxygen \\ atoms}} \frac{1 - \left(\frac{r_{Mg\cdots O_i}}{3}\right)^{12}}{1 - \left(\frac{r_{Mg\cdots O_i}}{3}\right)^{24}} \qquad (S1)$$

where $r_{Mg\cdots O_i}$ is the distance between $Mg^{2+}$ and oxygen atom $i$. The oxygen atoms of ADP are included in the coordination number, because the number of oxygen atoms interacting with $Mg^{2+}$ can change between 1 and 2. Past work showed that the coordination number of the cation ($Mg^{2+}$) is a more effective CV for the transition of CIP to SSIP than the coordination number of the anion(*27*, *29*, *30*, *89*, *90*), though this can depend on the identity of the ions.

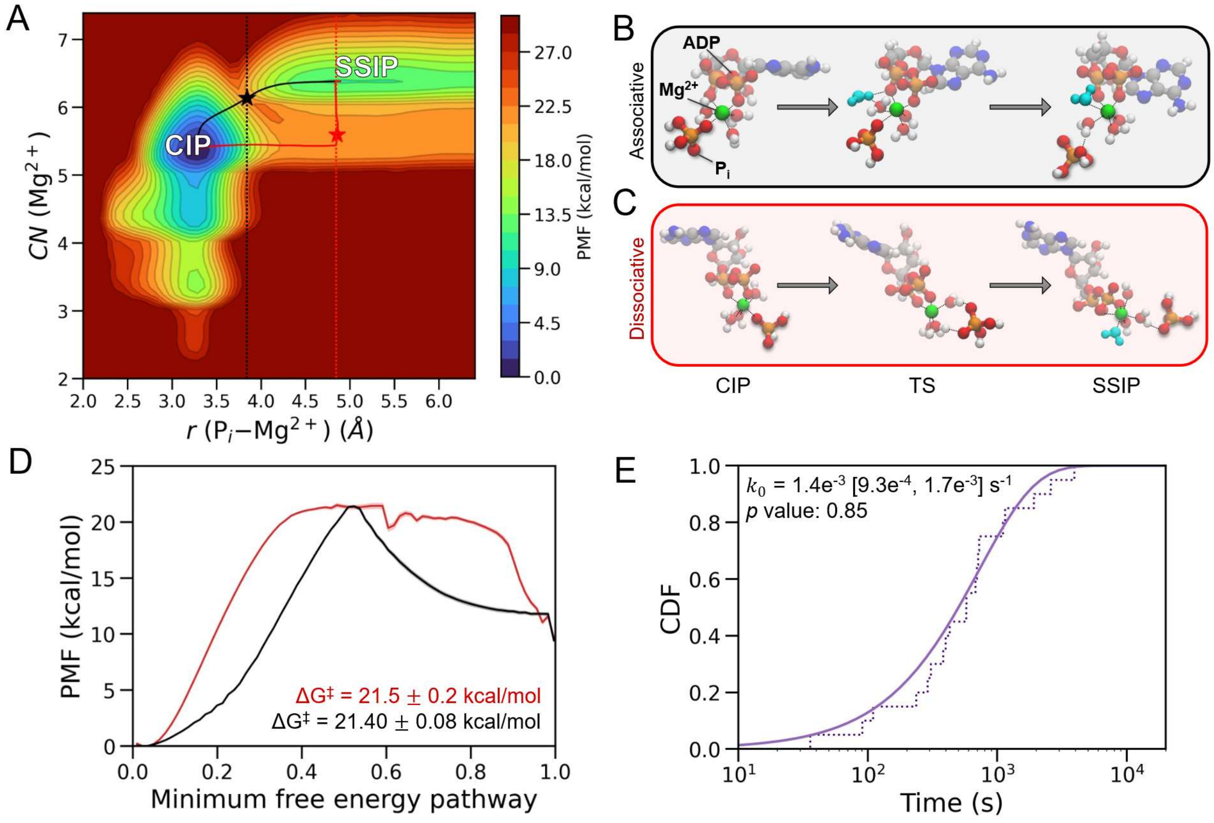


**Figure S2**. (A) 2-dimensional potential of mean force (PMF) for the transition from CIP to SSIP for ADP-$Mg^{2+}$-$P_i$ in TIP3P water. The CVs describing the transition are the distance between $Mg^{2+}$ and the center-of-mass of $P_i$, $r$ ($P_i$-$Mg^{2+}$), and the coordination number, $CN$ ($Mg^{2+}$), of oxygen atoms of water and ADP coordinated with $Mg^{2+}$. The red and black lines denote the minimum free energy pathways connecting the CIP and SSIP states. The red and black stars indicate the positions of the transition states. (B) The mechanism of the associative pathway for ion pair dissociation. Prior to elongation of the ion-ion distance, a water molecule (cyan) begins to insert (O···$Mg^{2+}$ distance: 3.37 Å) into the first solvation shell of $Mg^{2+}$. (C) The mechanism of the dissociative pathway for ion pair dissociation. The ions separate before the octahedral coordination of $Mg^{2+}$ is restored by a new water molecule (cyan). (D) The PMF along the minimum free energy pathways with the colors corresponding to the pathways in panel A. The $x$-axis represents the progression along the MFEP (unitless). Zero indicates the CIP state and one indicates the SSIP state. (E) The empirical cumulative distribution function (CDF) which depicts 20 rescaled transition times calculated using the method of Tiwary and Parrinello.(*34*) Each step (dotted line) is a time when one of the simulations finished by reaching the SSIP state. The fitted CDF yields a rate constant of $1.4e^{-3}$ [$9.3e^{-4}$, $1.7e^{-3}$] $s^{-1}$ and passes the Kolmogorov-Smirnov (KS) test with a $p$ value of 0.85. The uncertainty in the rate is estimated by bootstrapping.

The 2-dimensional PMF has two minima (Fig. S2A) representing the CIP state ($x \approx 3.3$ Å, $y \approx 5.4$) with $Mg^{2+}$ coordinated with $P_i$ and the SSIP state ($x \approx 4.9$ Å, $y \approx 6.4$) with $Mg^{2+}$ separated from $P_i$ by a water molecule. Note that the minima along the vertical axis do not align exactly at $CN$ ($Mg^{2+}$) = 5 for the CIP and $CN$ ($Mg^{2+}$) = 6 for the SSIP states, because $CN$ ($Mg^{2+}$) is a continuous function (Equation S1). In the CIP state (Fig. S1), $Mg^{2+}$ directly coordinates an oxygen atom of $P_i$ along with 5 other oxygen atoms of water/ADP. In the SSIP state (Fig. S1), $Mg^{2+}$ is separated from $P_i$ by a water bridge and is coordinated directly with 6 oxygen atoms of water/ADP. $Mg^{2+}$ strongly prefers an octahedral binding arrangement (either 5 interactions with water/ADP and 1 with $P_i$ or 6 interactions with water/ADP, see Fig. S2B-C) which is evidenced by the deep minimum of the CIP along the $CN$ ($Mg^{2+}$) degree of freedom. Note that the oxygen atom of $P_i$ is not counted in the $CN$ ($Mg^{2+}$) function to reflect the change in binding from CIP to SSIP (oxygen atom of $P_i$ vs. water molecule). The strong preference for octahedral binding agrees with experimental measurements of the coordination number of $Mg^{2+}$ in water.(*91–93*)

We delineated two minimum free energy pathways (MFEPs) from CIP to SSIP using the string method on the 2D PMF.(*94*) These two pathways represent associative (black line) and dissociative (red line) mechanisms (Fig. S2B-C), which differ in the order that $P_i$ dissociates from $Mg^{2+}$ and the new water molecule inserts into the first shell of $Mg^{2+}$. In the associative mechanism (black line), a water molecule (cyan) inserts between the first and second solvation shells of $Mg^{2+}$, after which $P_i$ dissociates from $Mg^{2+}$ (Fig. S2B). In the dissociative mechanism (red line), $P_i$ dissociates from $Mg^{2+}$, leaving $Mg^{2+}$ undercoordinated. A water molecule from the second solvation shell (cyan) reestablishes the stable octahedral coordination in the first solvation shell of $Mg^{2+}$ (Fig. S2C).

We integrated over the points ($r$, $CN$) in the direction orthogonal to the MFEPs to obtain the PMFs for the two minimum free energy pathways (Fig. S2D). The shapes of the two curves differ, yet the heights of the free energy barriers ($\Delta G^{\ddagger} \approx 21 - 22$ kcal/mol) along the two pathways (Fig. S2D) are thermodynamically indistinguishable within uncertainty.

The flatter transition state of the PMF for the dissociative mechanism (red curve) indicates a longer-lived transition state. Conversely, the sharper transition state of the PMF for the associative mechanism (black curve) indicates a transient transition state with few examples within the resolution of the saved simulation frames. Experimentally, a dissociative mechanism is thought to predominate for water exchange in the first solvation shell of $Mg^{2+}$ based on measurements of the activation volume,(*95*) though similar experiments have not been performed for $Mg^{2+}$-$P_i$.

Experimental measurements of the dissociation of $Mg^{2+}$ from DNA,(*96*) 5s rRNA,(*97*) and acetyl-phosphate(*41*) yielded estimated activation barriers of 12.7-13.3 kcal/mol. Acetyl-phosphate ($\Delta G^{\ddagger}_{exp.} = 13.1 \pm 0.2$ kcal/mol at 298 K) offers the closest comparison to the ADP-$Mg^{2+}$-$P_i$ ion pair. That said, the barrier of ~21.4 kcal/mol obtained with the CHARMM36m potential is likely overestimated by several kcal/mol which agrees with observations in the literature that nonpolarizable interaction potentials overstabilize CIPs.(*42–45*)

Using a reparameterization for $Mg^{2+}$ defined to match the water exchange in the first solvation shell,(*98*) the barrier for $P_i$ dissociation from ADP-$Mg^{2+}$ is notably lower ($\Delta G^{\ddagger}$ =17.9 kcal/mol) for the associative mechanism and only modestly lower ($\Delta G^{\ddagger} =$ 20.5 kcal/mol) for the dissociative mechanism (Fig. S3).

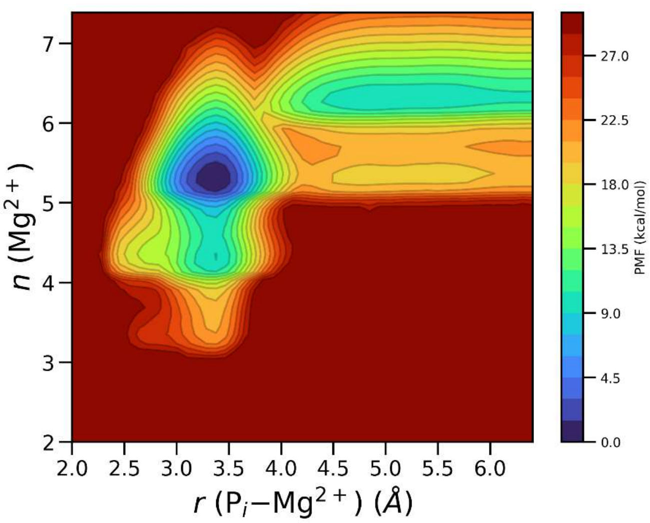


**Figure S3.** 2-dimensional potential of mean force (PMF) for the transition from CIP to SSIP for ADP-$Mg^{2+}$-$P_i$ in TIP3P water using the modified CHARMM potential. The axes represent the CVs which are the same as those used to obtain the PMF in Fig. S2A.

We used the polarizable AMOEBABIO18 force field(*99*, *100*) to investigate the role of polarizability on the absolute barrier height of the dissociation of $P_i$ from ADP-$Mg^{2+}$. The parameters for ADP were obtained from Walker et al.(*101*) The parameters for $P_i$ were obtained from Kumar et al.(*102*) The parameters for ADP, $Mg^{2+}$, and $P_i$ were modified based on the HFC23 revision.(*103*) Short-range interactions were computed in real space up to a cutoff of 9 Å. We used an Ewald summation to compute long-range interactions in reciprocal space. The simulations with the polarizable interaction potential were performed using the GPU implementation(*104*) of the Tinker-HP software(*105*) patched with PLUMED 2.7.(*84*)

Fig. S4 shows the PMFs obtained with the polarizable AMOEBABIO18(*99*–*102*) and AMOEBABIO18-HFC23(*103*) force fields. The polarizable interaction potentials exhibit lower activation barriers, though the barrier height depends sensitively on the parametrization.

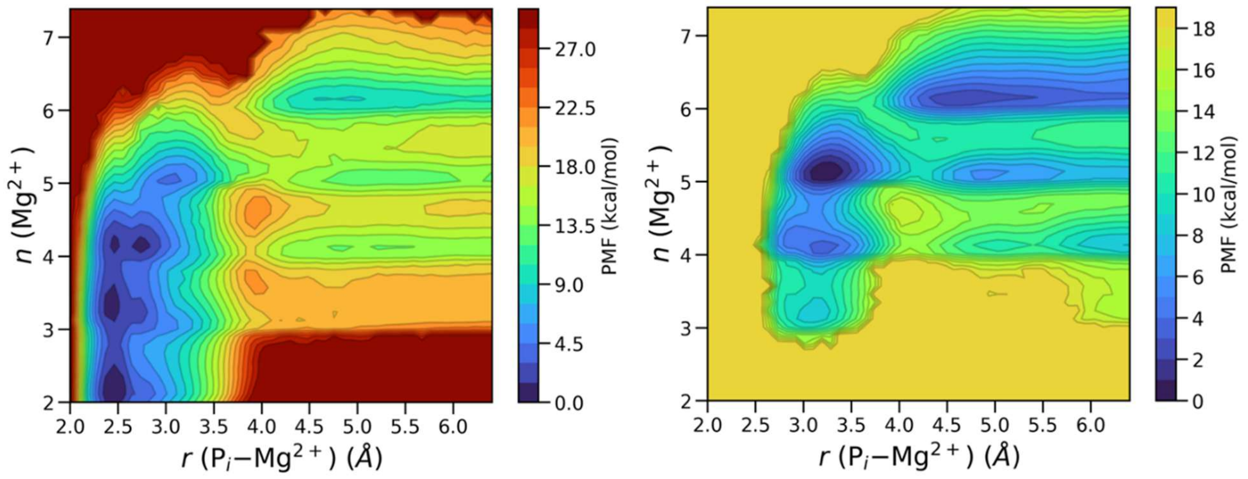


**Figure S4.** 2-dimensional potential of mean force (PMF) for the transition from CIP to SSIP for ADP-$Mg^{2+}$-$P_i$ in TIP3P water using the AMOEBABIO18 (left) and AMOEBABIO18-HFC23 (right) potentials. The axes represent the CVs which are the same as those used to obtain the PMF in Fig. S2A.

The free energy wells for the CIP and SSIP states overlap along $r$ ($P_i$-$Mg^{2+}$) on the $x$-axis (Fig. S2A) which demonstrates the importance of choosing effective CVs. The transition state in the associative mechanism (black star) overlaps on the $x$-axis with lower energy conformations of the CIP state whereas the transition state for the dissociative mechanism (red star) overlaps with lower energy conformations of the SSIP state, indicated by dotted lines. Fig. S5 shows 1-dimensional cuts from the 2D PMF to illustrate this effect more explicitly.

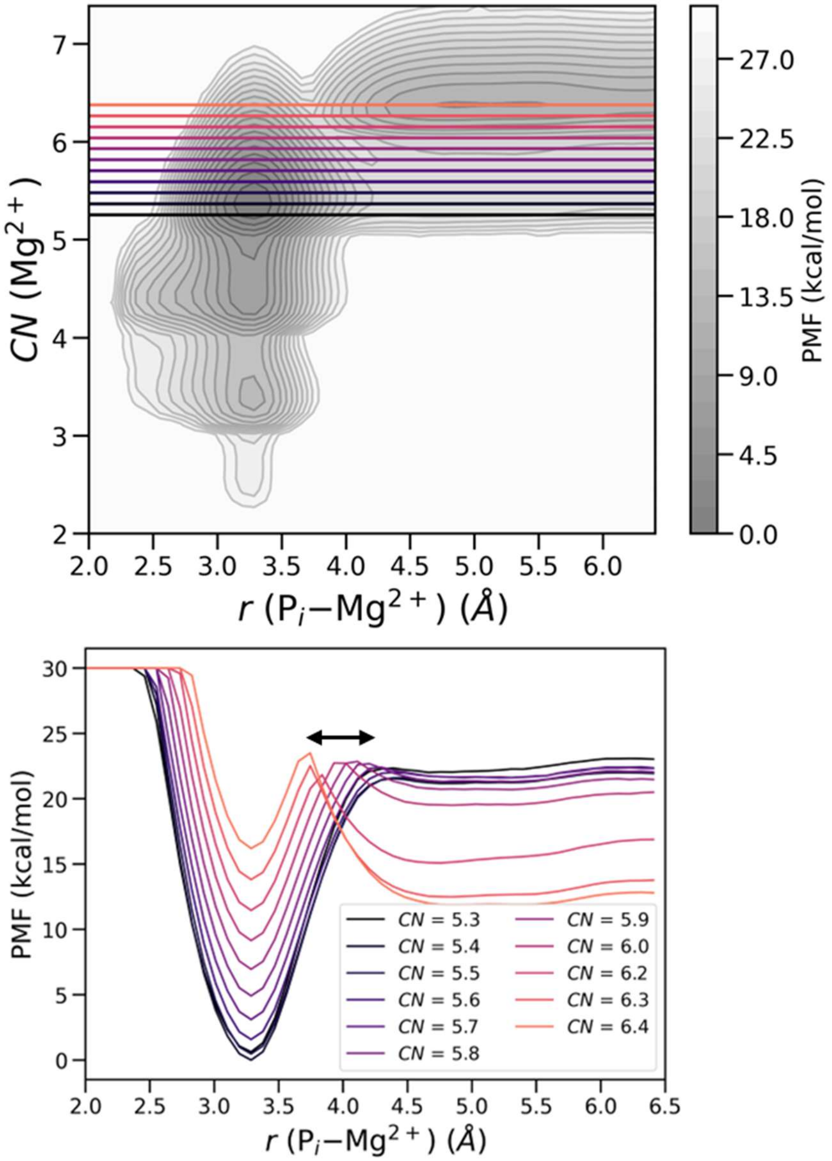


**Figure S5.** The 2D PMF with the CHARMM36m potential (top) in grayscale with the colored horizontal lines corresponding to the 1D PMF cuts at those $CN$ ($Mg^{2+}$) values (bottom). The arrow denotes the range of barrier height positions, showing the overlap of the CIP and SSIP wells along the $x$-axis.

Consideration of only the $r$ ($P_i$-$Mg^{2+}$) degree of freedom (Fig. S6) yields a barrier of ~17.3 kcal/mol which is ~ 4 kcal/mol lower than the barrier estimated using two CVs. The insufficiency of describing the PMF with $r$ ($P_i$-$Mg^{2+}$) alone has been noted for other ion pairs(*27*, *28*, *30*, *42*) but is especially prominent due to the strong preference of $Mg^{2+}$ for octahedral coordination.

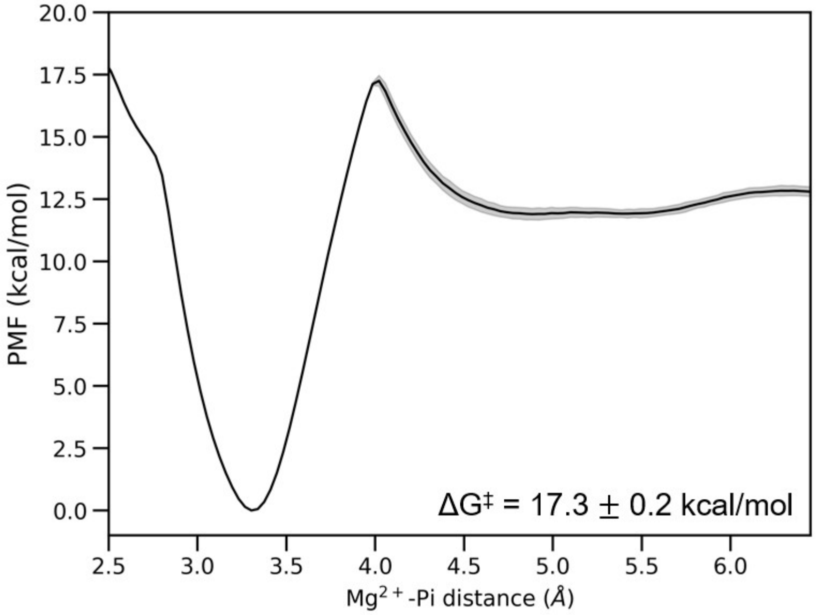


**Figure S6.** The potential of mean force (PMF) along the $Mg^{2+}$-$P_i$ distance. This apparent barrier is ~4 kcal/mol lower than when the coordination number of $Mg^{2+}$ is included as a CV (Figure 2).

### Section 2: Kinetics of the transition of the contact ion pair (CIP) to solvent-separated ion pair (SSIP) for ADP-$Mg^{2+}$-$P_i$ in water

We estimated the rate constant of transition from CIP to SSIP in two ways. Using transition state theory (TST, Equation 1) and the heights of the energy barriers, we estimated the rate constant for both pathways to be $k_{TST} = 0.0007$ $s^{-1}$ with a lower bound of 0.0005 $s^{-1}$ and upper bound of 0.0010 $s^{-1}$] which reflect the uncertainties in the barrier heights (Fig. S2D).

We also estimated the rate constant directly using the method of Tiwary and Parrinello.(*34*) Fig. S2E shows the empirical cumulative distribution function (CDF) of the rescaled transition times from 20 independent simulations of the transition of CIP to SSIP. The time-acceleration factor $\alpha$ is estimated as the time integral of the bias (see Methods for details), which were deposited along $r$ ($P_i$-$Mg^{2+}$) during MD simulations which we initiated from the CIP state. Each simulation ended once the system transitioned from the CIP to SSIP. Fitting the empirical CDF yields the estimated rate constant $k_0$ of 0.0014 $s^{-1}$ for the transition from CIP to SSIP with an uncertainty range of 0.0009 to 0.0017 obtained by bootstrapping.

Note that the challenge with CVs discussed above is not an issue with this approach, because it only samples and applies bias to the CIP state, while the PMF calculation requires sampling both the CIP and SSIP and the use of $CN$ ($Mg^{2+}$) to fully distinguish the states.

We used the Kolmogorov-Smirnov (KS) test to determine whether the transition times observed during simulations are from a Poissonian distribution ($CDF = 1 - e^{-k_0 t}$). The $p$ value of 0.85 is much larger than the minimum threshold of 0.05, indicating an excellent match to the Poissonian distribution (solid line in Fig. S2E).

The estimated rate of 0.0014 [0.0009, 0.0017] $s^{-1}$ agrees with the upper range of the estimate (0.0010 $s^{-1}$) obtained from transition state theory using the free energy barrier from the PMF (Fig. S2D). This shows that the pairwise distance CV is sufficient to promote transition from CIP to SSIP to obtain rescaled first passage times, whereas the 2nd CV (coordination number of $Mg^{2+}$) is necessary to separate the CIP, SSIP, and transition states in the PMF. The correspondence between the rate constant and the barrier height obtained from separate approaches indicates that the CVs effectively describe the transition from CIP to SSIP.

**Additional computational details for Sections 1 and 2:**

**PMF calculations.** The unbiased histogram, $P(r, CN)$, representing the probability that the system exists in state $(r, CN)$ was obtained using the reweighting factors (CALC_RCT) calculated in PLUMED.(*106*) The PMF in Fig. S2D was computed from the unbiased histogram as

$$PMF(r, CN) = -k_B T ln\, P(r, CN) \qquad (S2)$$

For the 1D case, we integrated over the $CN$ degree of freedom to obtain $PMF(r)$. The minimum free energy paths (MFEPs) were obtained using the string method.(*94*) Each point defined by $r$ and $CN$ was mapped to the nearest point on the MFEP. In other words, the points were mapped in a way such that the line connecting each point and the MFEP is orthogonal to the tangent of the MFEP at that point. Subsequently, we integrated over the orthogonal direction to obtain the PMF along the MFEP. The uncertainty was obtained by block analysis, using the last ¾ of the trajectory to estimate the average and standard error of the MFEPs over 3 blocks. The bias height was set at 1.0 kJ/mol, width at 0.05 Å and 0.05, and bias factor to 10. Bias was deposited every 5 ps. An upper wall with a force constant of 200 kJ/$Å^2$ was placed at $r$ ($P_i$-$Mg^{2+}$) = 6.5 Å to sample the CIP to SSIP (and reverse transition) several times.

**Transition state theory (TST).** TST assumes that reactants must proceed through an activated transition state before relaxing to the product state in a single elementary step. Equation S3 provides a connection between the activation barrier height ($\Delta G^{\ddagger}$), temperature ($T$), and first-order rate constant ($k_{TST}$):

$$k_{TST} = \frac{\kappa\omega_0}{2\pi} e^{-\Delta G^{\ddagger}/k_B T} \quad \text{(S3)}$$

$k_B$ is the Boltzmann constant and $\kappa$ is the transmission coefficient which accounts for barrier recrossing. The transmission coefficient is often assumed to be unity, but past work has established that recrossing cannot be eliminated entirely for ion pair dissociation.(*29*) $\omega_0$ is the vibrational frequency along the reaction coordinate which is estimated as $\frac{k_B T}{h}$ in the Eyring equation.(*107*, *108*) For a more accurate estimate, we estimate $\omega_0 = \sqrt{\frac{\langle \dot{q}^2 \rangle}{\langle q^2 \rangle}}$ for the reactant (CIP) state where $q$ is the reaction coordinate. We note the existence of additional corrections for the motion of the reaction coordinate for the reactants and for the curvilinear nature of the reaction coordinate (via the inverse of the effective mass of the reaction coordinate),(*109*, *110*) though we do not explore these in the present work. While reactions/processes can pass through several intermediates, each with their own free energy barriers, the largest barrier in the process (the rate-limiting step) generally determines the rate of the entire process.

### Section 3: Ion pair dissociation in the phosphate cavity of actin filaments

Figure S7 shows an example of the CIP and SSIP states in the phosphate cavity of the terminal barbed end subunit. This can be compared to the examples in the interior subunit in Figure 1C. Notably, the ion pair in the barbed end subunit have a more extensive hydrogen bond network in the first solvation shell (6-7 molecules in Fig. S7) compared to the interior subunit (Fig. 1B).

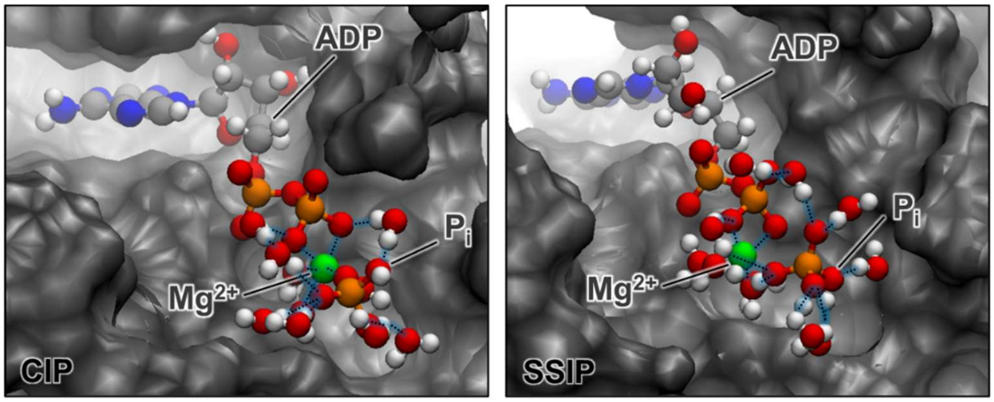


**Figure S7.** Snapshots of the transition from the CIP state (left panel) to SSIP state (right panel) in the phosphate cavity of the barbed end subunit. The dark gray is the surface of the protein computed with VMD with a probe radius of 1.4 Å. Only water molecules with oxygen atoms within 4.5 Å of the phosphorus atom of $P_i$ or within 2.5 Å of the magnesium atom are shown.

The volumes of the phosphate cavities differ for the pointed end, barbed end, and interior subunits, which we illustrate with ribbon diagrams with space-filling rendering of the phosphate cavities for each subunit. The frames used for the visualization were selected to show the volume near the minimum (left) average (middle), and maximum (right) for that subunit (Fig. S8).

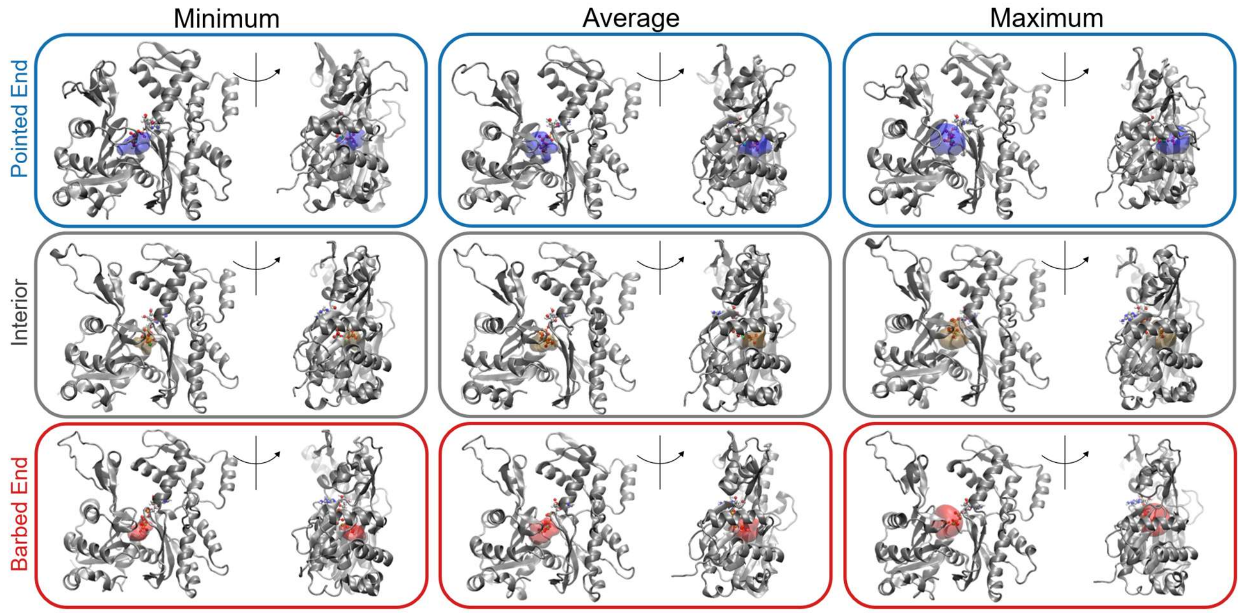


**Figure S8.** Comparison of phosphate cavity volumes for the filament ends and the interior. The frames shown were selected to show minimum (left), average (middle), and maximum (right) volumes over a simulation.

### Section 4: Hydrogen bond lifetime in the protein cavity near the CIP

We computed the continuous and intermittent hydrogen bond lifetimes (HBLs) for the hydrogen bonds between $Mg^{2+}$ and its 1st solvation shell, $P_i$ and its 1st solvation shell, the 1st and 2nd solvation shells of $Mg^{2+}$, and the 1st and 2nd solvation shells of $P_i$. We found it valuable to consider the HBLs by solvation shells, since proximity to the ions influences their dynamics. The water – water and water – $P_i$ hydrogen bonds were detected using the geometric definition of the hydrogen bond, which specifies an O···O distance ≤ 3.5 Å and an O-H···O angle ≥ 100°. A cutoff distance of $r$ ($Mg^{2+}\cdots O_w$) < 2.5 Å was used to define $Mg^{2+}$ – $H_2O$ bonds.

We used 30 simulations of 0.1 ns to obtain the HBLs. For the actin systems, the starting points for the simulations were obtained at 20 ns intervals from an unbiased simulation, spanning 600 ns of simulation time in total. This helped to sample across slower changes in the phosphate cavity size and hydrogen bond network. The frames were saved every 50 fs. Because we split the HBLs by solvation shell (and there are fewer molecules to average over), it was necessary to obtain results from numerous simulations to obtain reasonable statistics. The HBLs were computed using the following autocorrelation function,

$$C(\tau) = \frac{\langle h(0)h(\tau)\rangle}{\langle h(0)h(0)\rangle} \tag{S4}$$

where $h(t)$ is 1 if a hydrogen bond exists at time $t$ and 0 if not. The intermittent HBLs account for the reforming of hydrogen bonds at a time after the hydrogen bond is first broken. The continuous HBLs measure the lifetimes of a single hydrogen bond. Specifically, if $h(t) = 0$, then $h(\tau > t) = 0$.

**Continuous hydrogen bond lifetimes**

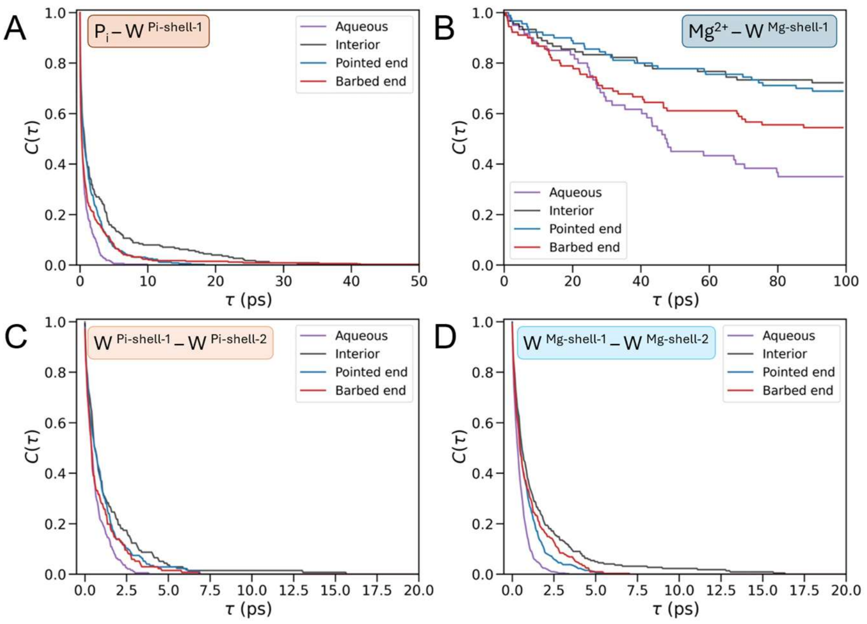


**Figure S9.** The continuous HBLs for (A) $P_i$ with its first solvation shell, (B) $Mg^{2+}$ with its first solvation shell, (C) between waters in the first and second solvation shells of $P_i$, and (D)between waters in the first and second solvation shells of $Mg^{2+}$. The HBLs were computed for simulations of the CIP in water (purple), in the interior subunit (dark grey), in the pointed end (blue), and barbed end (red). The insets portray graphical representations of the hydrogen bond network.

Figure S9 depicts the continuous HBLs for the hydrogen bonds between $P_i$ and water molecules in its 1st solvation shell, $Mg^{2+}$ and water molecules in its 1st solvation shell, the water molecules in the 1st and 2nd solvation shells of $Mg^{2+}$, and the water molecules in the 1st and 2nd solvation shells of $P_i$. As expected, the hydrogen bond dynamics are very slow near $Mg^{2+}$ (and $P_i$ to a lesser extent). Notably, the HBLs are much longer in water (note the different time scales) than in actin filament phosphate cavities. Amongst the actin filament phosphate cavities, confined protein cavities slow the hydrogen bond dynamics with HBLs longer in interior subunits than subunits at the ends of the filaments and in water.

**Intermittent hydrogen bond lifetimes**

The intermittent HBLs provide insights into the persistence of the hydrogen bond network (Figure S10). As expected, the molecules in the first solvation shell of $Mg^{2+}$ do not change over the course of any of the short simulations (Fig. S10B). In water (purple), very few water molecules in the first solvation shell of $P_i$ at time $\tau = 0$ remain at $\tau = 0.1$ ns (Fig. S10A). In the protein, 40-70% of the water molecules in the first solvation shell of $P_i$ at $\tau = 0$ remain at $\tau = 0.1$ ns (Fig. S10A). Similar values are observed for the 2nd shell HBLs showing the persistent hydrogen bond network in the confined protein cavity (Fig. S10C).

In general, the hydrogen bond network rearranges more quickly around the $Mg^{2+}$ ion in the barbed end subunit than in the interior and pointed end subunits. Conversely, it arranges more quickly around $P_i$ in the pointed end (than in the interior and barbed end subunits).

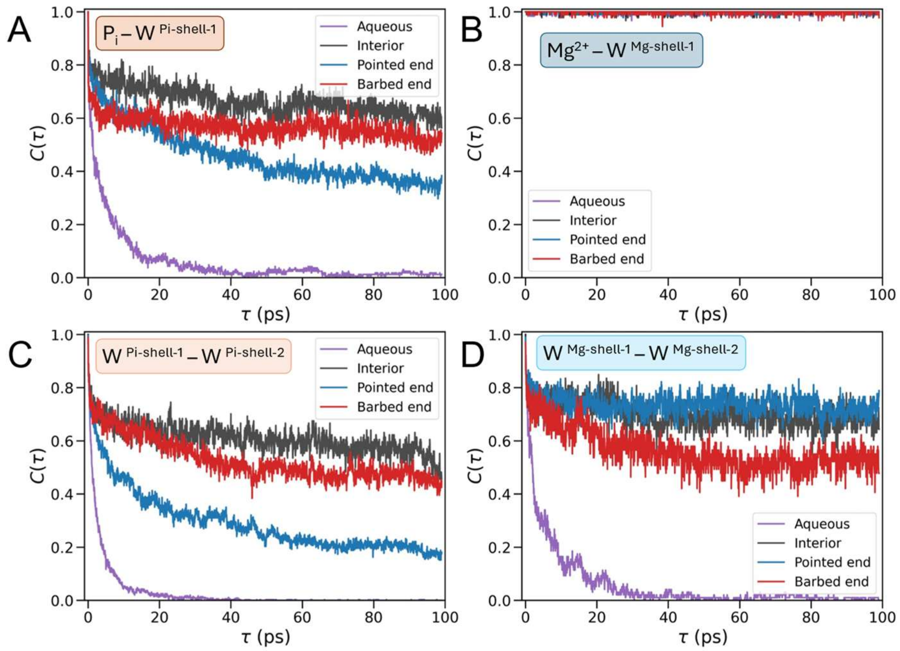


**Figure S10.** The intermittent HBLs for (A) $P_i$ with its first solvation shell, (B) $Mg^{2+}$ with its first solvation shell, (C) the first and second solvation shells of $P_i$, and (D) the first and second solvation shells of $Mg^{2+}$. The HBLs are computed for simulations of the CIP in water (purple), and in actin subunits of the interior (dark grey), pointed end (blue), and barbed end (red) of the filament.

### Section 5: Relationship between protein conformations and phosphate cavity volumes

We measured contacts between $P_i$ and amino acid residues in the phosphate cavity across ~1.6 μs of unbiased simulation (Table S1). $P_i$ has more contacts with protein in interior subunits than terminal end subunits. Specifically, in the interior subunit, close interactions of $P_i$ several residues of the P2 loop (residues 154-161) and the P1 loop (residues 11-14) exclude water molecules near the CIP.

**Table S1.** The percentage of MD frames in which the heavy atoms of amino acids contact the heavy atoms of $P_i$ (within a cutoff of 4.5Å) during ~1.6 μs of unbiased simulation time. Note that amino acids in contact with $P_i$ in less than 25% of the MD frames are not included.

| | **Interior** | **Barbed End** | **Pointed end** |
|---|---|---|---|
| **ASP11** | N/A | 50% | N/A |
| **ASN12** | N/A | 60% | 48% |
| **GLY13** | 35% | 60% | 46% |
| **SER14** | 78% | 50% | 88% |
| **GLY74** | 26% | 32% | 45% |
| **ALA108** | 33% | N/A | 90% |
| **GLN137** | 74% | 93% | 78% |
| **ASP154** | 87% | 72% | N/A |
| **SER155** | 73% | N/A | N/A |
| **GLY156** | 98% | N/A | N/A |
| **ASP157** | 88% | N/A | N/A |
| **GLY158** | 87% | N/A | 48% |
| **VAL159** | 94% | N/A | 50% |
| **HIS161** | 64% | N/A | 52% |
| **VAL339** | N/A | 45% | N/A |

**Distributions of normal modes at different ranges of phosphate cavity volumes**

We report distributions of normal modes for 5 subunits at four ranges of phosphate cavity volumes: 0-50 Å$^3$, 50-100 Å$^3$, 100-150 Å$^3$, and 150-200 Å$^3$ (Fig. S11). In general, the subunits at the ends are distinct from the interior subunits, even at similar volumes. For subunits B and P, we observe a shift in the scissors and dihedral angles distributions for larger volumes.

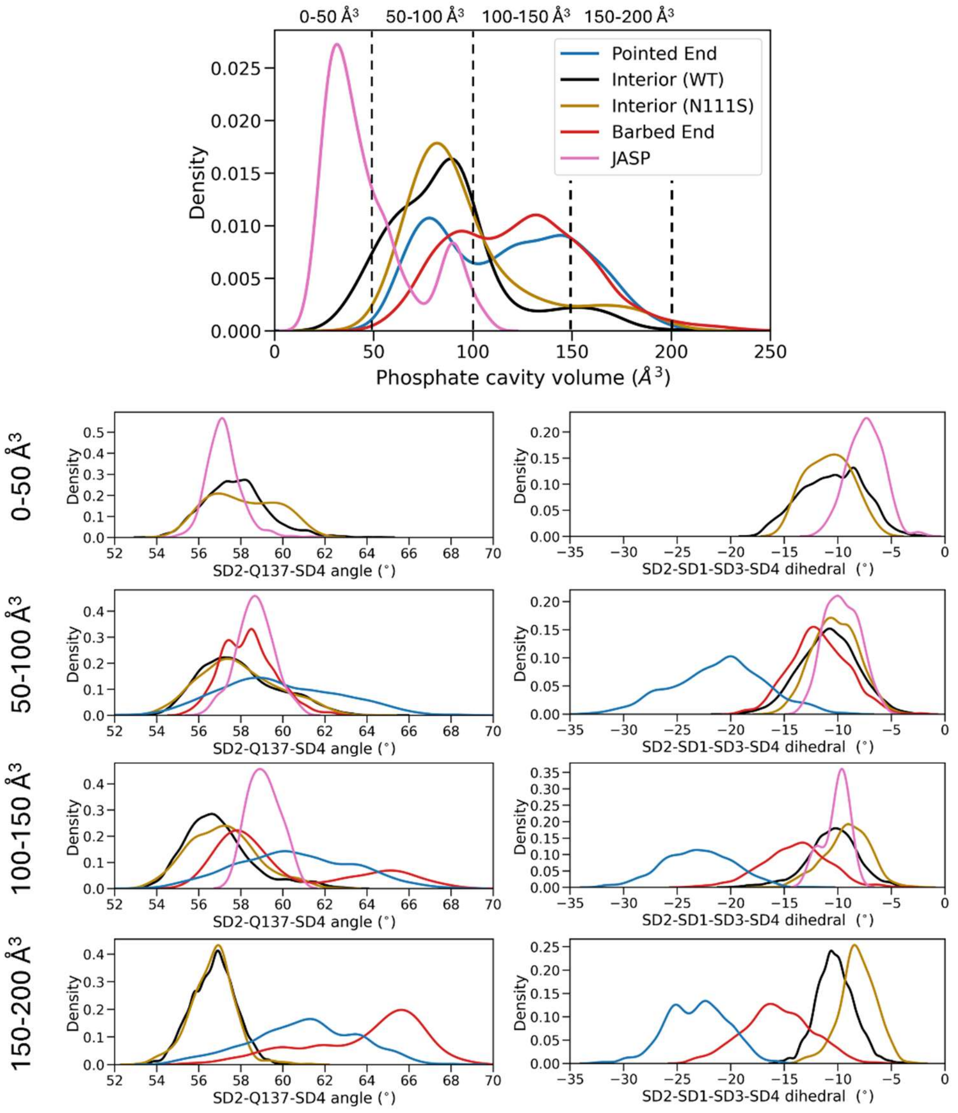


**Figure S11**. (Top graph) Distributions of phosphate cavity volumes during unbiased simulations of 5 subunits. The dashed lines highlight the four groups of volume ranges. (Bottom) The distributions of (left) scissors angles (SD2-Q137-SD4) and (right) subdomain dihedral across the four ranges of phosphate cavity volumes.

**Relation of scissors and dihedral angles to volumes of the phosphate cavity**

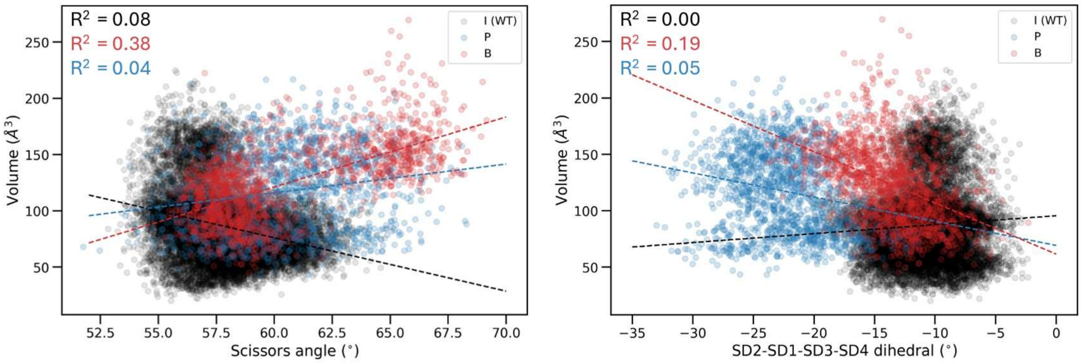


**Figure S12.** Correlation plots between the scissors (SD2-Q137-SD4) and SD2-SD1-SD3-SD4 dihedral angles and the phosphate cavity volume plotted every 1 ns across ~1.6 µs of unbiased simulation time. The dashed lines are the lines of best fit.

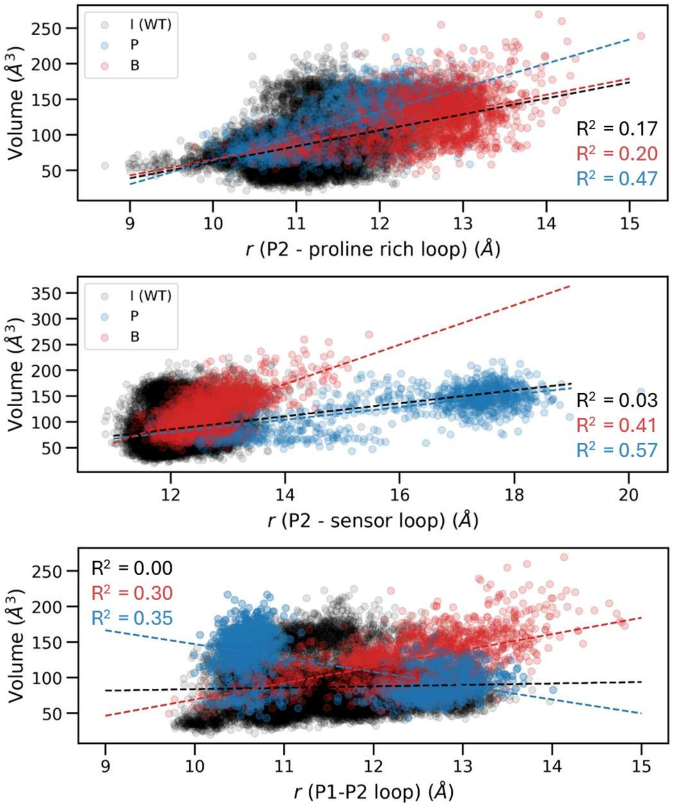


**Figure S13.** Correlations between the distances between the P2 – proline rich loop (top), P2 – sensor loop (middle), and P1 – P2 loop (bottom) and phosphate cavity volumes. The dashed lines represent the line of best fit.

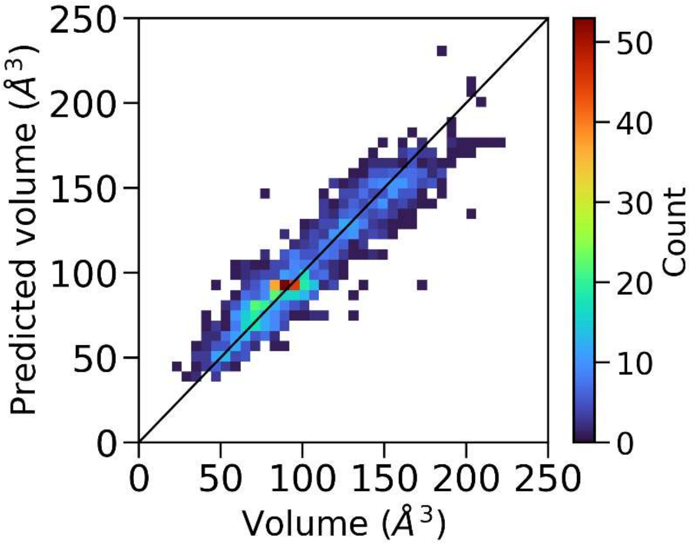


**Figure S14.** Correlation between volumes predicted by a Random Forest Regression model on the *y*-axis and volumes of the phosphate cavity on the *x*-axis at each frame of the test set of ~ 1,200 frames. The non-linear regression model was trained using the COM-to-COM distances between the P1, P2, sensor, and proline rich loops, the scissors and dihedral angles, and distances between the phosphorus atom of $P_i$ and the nearest heavy atom of residues 11-14, 74, 108, 137, 154-161 at ~6,300 MD frames. The mean absolute error (MAE) of the model on the test set is 10 $Å^3$ and the mean absolute percent error (MAPE) is 11%.

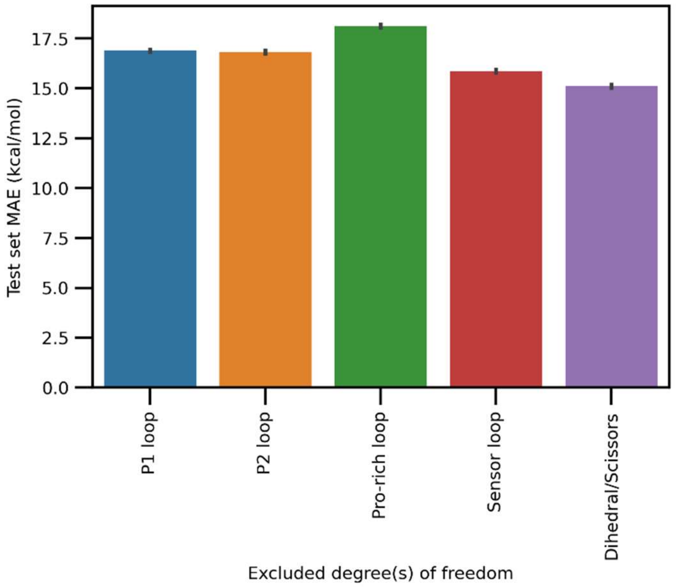


**Figure S15.** Mean absolute error (MAE) on the test set of Random Forest models trained with certain degrees of freedom (x-axis) excluded from the dataset. The error bars show the standard deviation across models trained on different subsets of the dataset. This analysis shows that the proline-rich loop and P1 and P2 loops are the most important descriptors as expected given that they directly border the phosphate cavity. A caveat of this analysis is the weak to moderate correlation between some of these variables (Fig. 3F-H).

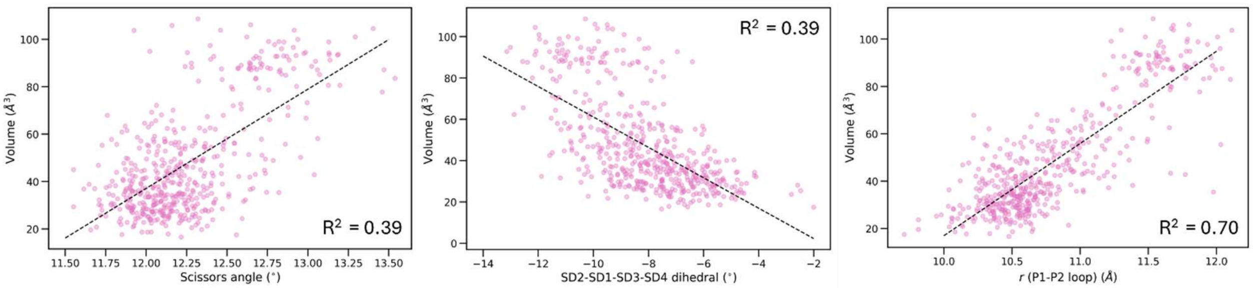


**Figure S16.** Correlations between the phosphate cavity volume (*y*-axis) in subunits with jasplakinolide bound and the scissors angle (left), SD2-SD1-SD3-SD4 dihedral angle (middle), and COM-to-COM distances between the P1 and P2 loops (right). The dashed black lines show the least squares fits.

**Comparison of root mean squared fluctuations residues in three subunits**

The root mean squared fluctuation (RMSF) for each residue in the subunits were computed for the terminal barbed end, pointed end, and interior subunits (Figure S17A). The D-loop of the pointed end subunit fluctuates the most, since SD2 and SD4 are unbound (Fig. S17C). The D-loop of the barbed end subunit fluctuates slightly more than interior subunits. The H-plug region of the barbed end subunit (residues 263-273, Fig. S17B) fluctuates more than other subunits, because the lateral contacts between B and B-1 can break and reform. The two α-helices of SD4 (nearest to SD2) of the terminal subunits fluctuate more than the interior subunit (residues 180-220). Residues 320-328 of SD3 and the region connecting SD1-SD3 (residues 333 to 340) at the front of the barbed end subunit fluctuate more than other subunits and help explain why $P_i$ egressed near residues 333 to 340 in some simulations of the barbed end subunit.

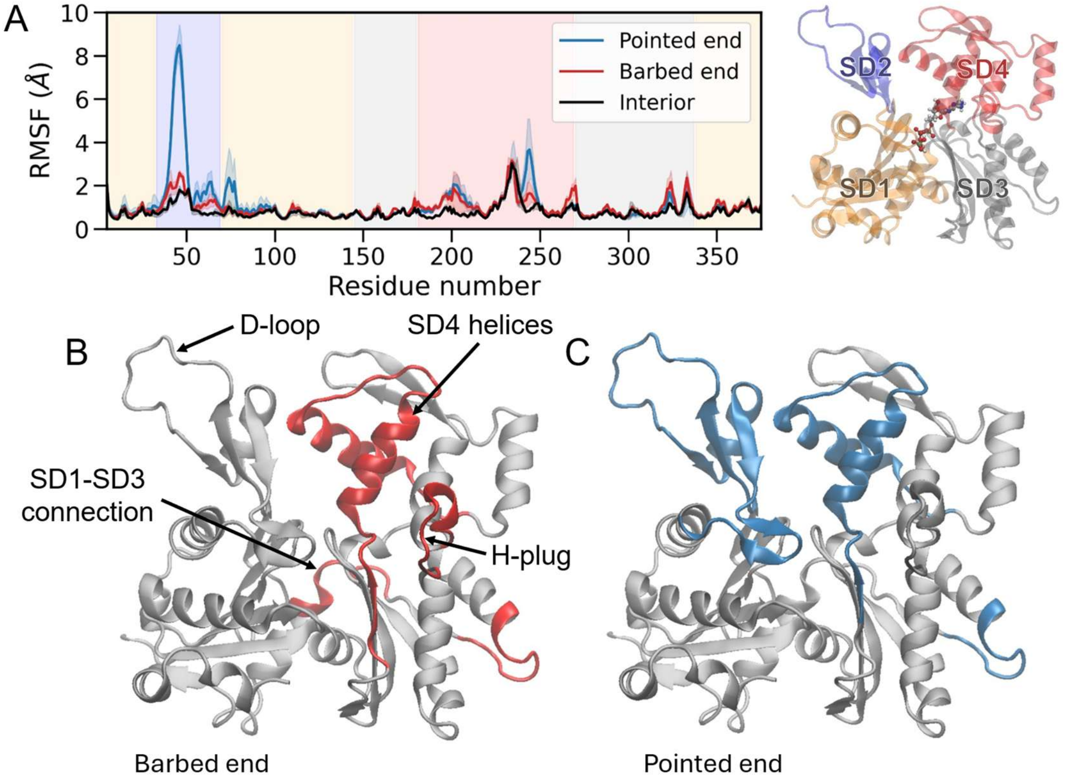


**Figure S17.** (A) Root mean squared fluctuation (RMSF) for each residue of interior (black), pointed end (blue), and barbed end subunits (red). The subdomains (SD) are shaded with different colors in the right image. Colored regions show sections of the barbed end (B) and pointed end (C) subunits that fluctuate more than the interior subunit.

**Section 6: Comparison of conformation of subunit P-1 to subunit P**

Subunit P-1 had a distribution of dihedral angles slightly more twisted than interior subunits, similar to barbed end subunits (Fig. S18). The scissors angles of subunit P-1 were notably smaller than all other subunits measured, likely due to lateral interactions between its D-loop of SD4 with subunit P.(*12*, *25*) The distances S14 to G148, R183 to Y69, and D72 to G158 were smaller in subunit P-1 than subunit P (Fig. S19)

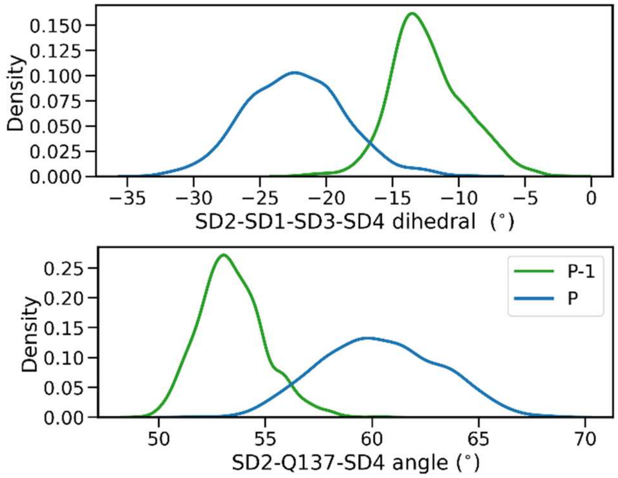


**Figure S18.** The distributions of SD2-SD1-SD3-SD4 dihedral (top) and scissors (SD2-Q137-SD4) angles (bottom) for subunits P (blue) and P-1 (green).

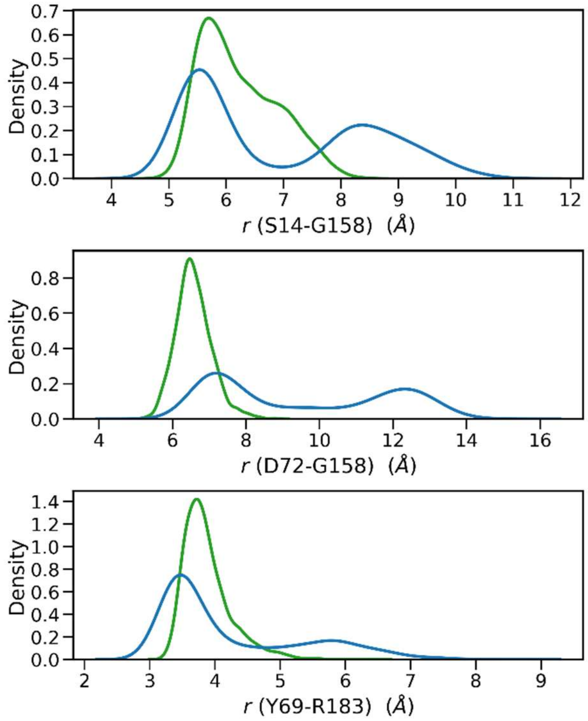


**Figure S19.** The distributions of residue-residue distances for the pathway above the sensor loop for subunits P (blue) and P-1 (green).

### Section 7: Effects of mutations on $P_i$ release

Interior subunits of N111S filaments released phosphate ≥ 15x faster than those of wild-type filaments in biochemical experiments.(*8*) This corresponds to a decrease of ~ 1.7 kcal/mol in the barrier height for release. In our simulations, the rates of CIP-to-SSIP transition in interior subunits of filaments composed of N111S mutant subunits were indistinguishable within uncertainties from filaments with wild type subunits (Fig. S20) and the cumulative distribution functions (dotted lines) overlapped. Thus, it could not distinguish how the N111S mutation impacts the kinetics of $P_i$ release.

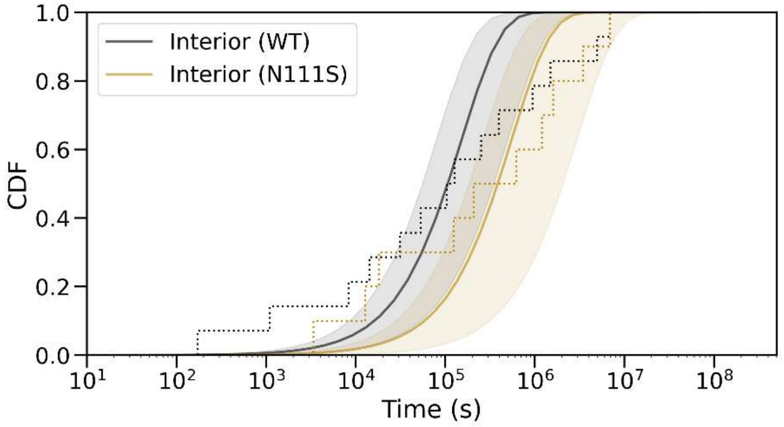


**Figure S20.** The empirical cumulative distribution functions (dotted lines) comparing the time courses for the CIP-to-SSIP transition in interior subunits of filaments with wild-type (black) and N111S (gold) subunits. The solid lines are best fits to the equation $(1 - e^{-k_0 t})$ where $k_0$ is the rate constant estimated from the fit. The shaded regions show the uncertainty in the fits estimated by bootstrapping.

Owing to a shorter sidechain S111 is separated further from R177 than N111 and the dynamics of the sensor and P2 loops differed from wild type subunits. During our simulations the volume of the phosphate cavity of N111S subunits was larger and more water molecules surrounded $P_i$ than in wild type subunits (gold line, Fig. S21). This larger phosphate cavity is indicative of a faster CIP-to-SSIP transition, but the simulation did not resolve the small difference in the barrier height.

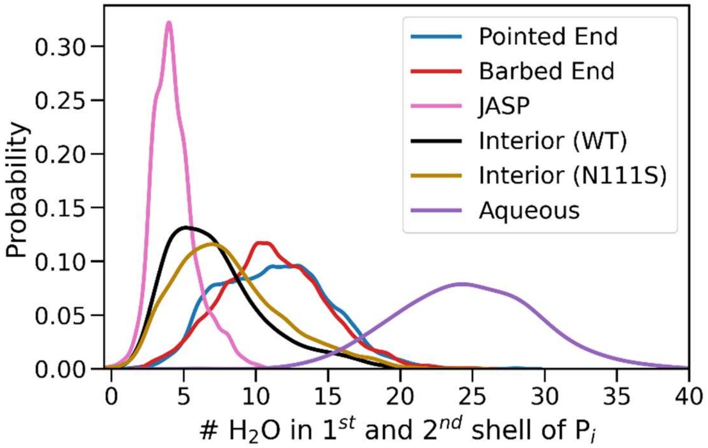


**Figure S21.** Distributions of the number of water molecules in the first and second solvation shell of Pi. Notably, the distribution corresponding to interior subunits of N111S filaments (gold) had a smaller population of states with low hydration (~ 3-7 water molecules) and a larger population of states with high hydration (~ 10-20 water molecules).

We also did not detect significant difference in the PMF for the egress from the SSIP state to the protein exterior (Fig. S22-S24). The pathways for release of phosphate from interior subunits of N111S filaments are similar to those for interior subunits of wild-type filaments. The simulations show a slight preference for the green pathway (3 of 5 simulations).

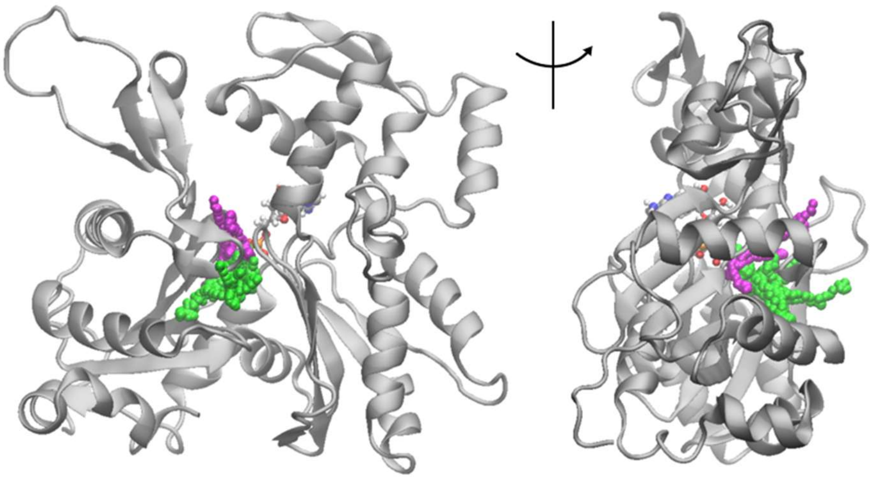

**Figure S22.** Ribbon diagrams of interior subunits with colored sphere showing the egress pathways from 5 independent WT-MetaD simulations of actin filaments with the N111S mutation. In 3 simulations, $P_i$ egressed through the back door route under the sensor loop (green) and, in 2 simulations, $P_i$ egressed through the back door route above the sensor loop (magenta).

**Mutations of residue R183 (distant from N111-R177 gate)**

Mutating R183 had a small effect on the rate of $P_i$ dissociation from interior subunits of actin filaments in experiments: only 1.8 faster for R183G and 2.9x faster for R183W.(*8*) These mutations eliminate the cation-$\pi$ interaction between R183-Y69. Loss of this interaction would likely influence the SD2-SD4 and P1-P2 loop distances which we show are determinants of the volume of the phosphate cavity. This may help to explain the slight decreases in barrier height (approximately 0.4 and 0.7 kcal/mol) for mutations distant from the N111-R177 gate.

### Section 8: Thermodynamics of $P_i$ release pathways from different subunits

We obtained potentials of mean force (PMF) from the egress simulations using the deposited bias to reweight the probability distribution along the collective variable, the $Mg^{2+}$-$P_i$ distance. These PMFs focus on barriers during egress from the SSIP state (4-6 Å on the *x*-axis of Fig. S23) to the protein exterior. The PMFs are shifted so that PMF = 0 kcal/mol once $P_i$ fully exited from the protein.

$P_i$ passes through the R177-N111 gate at $Mg^{2+}$-$P_i$ distances ranging from approximately 11-16 Å in the green pathway. There is not a substantial barrier in this region.

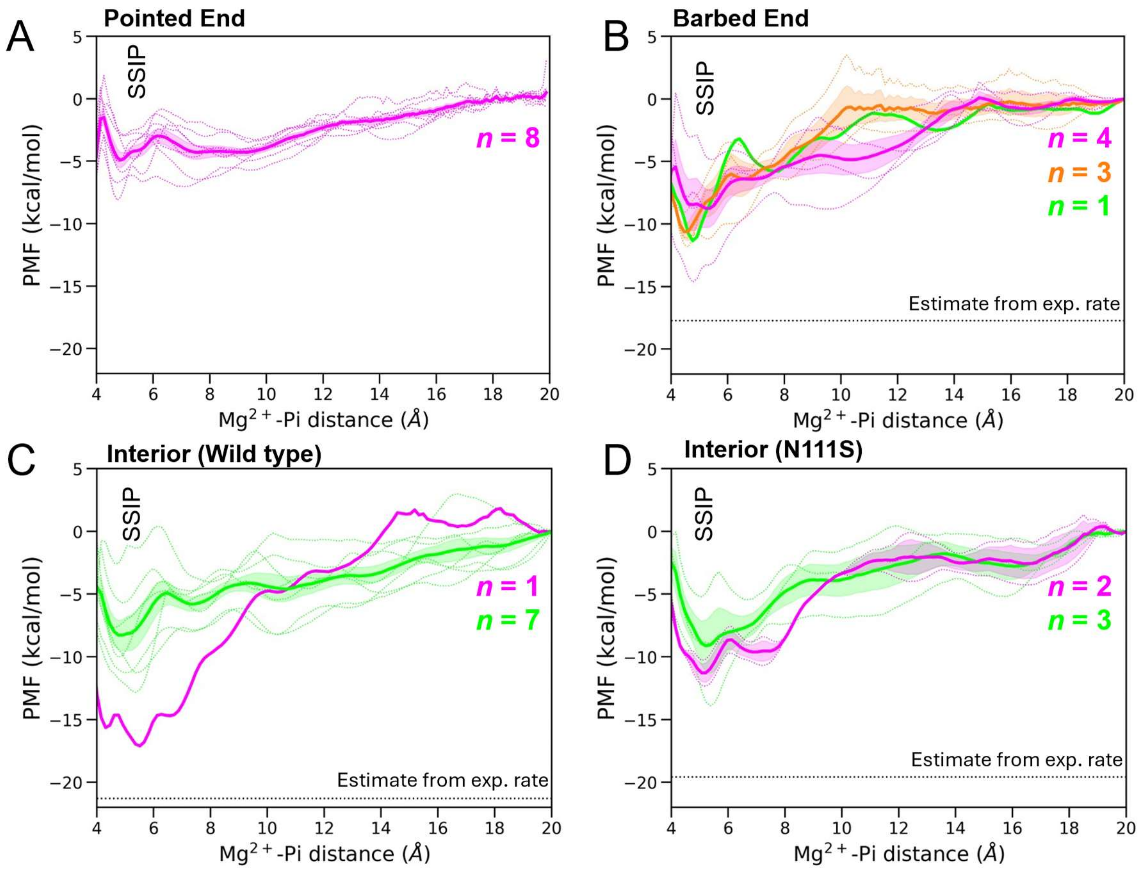


**Figure S23.** PMFs of $P_i$ egress from the SSIP state to the protein exterior for four subunits: (A) the pointed end; (B) barbed end; (C) interior (wild type); and (D) interior (N111S). The colors represent the category of egress pathway: green, back door below sensor loop; magenta, back door above sensor loop; and orange, front door. The traces represent the results from independent simulations. The dark line is the average, and the shaded regions show the standard error.

#### Cumulative distribution functions (CDF) for the transition of SSIP to the exterior of the protein

Using the method of Tiwary and Parinello,(*34*) we estimated rescaled passage times from the SSIP state to the exterior of the protein for the pointed end (blue), barbed end (red), interior (WT, black), and interior (N111S mutant, gold) in Fig. S24. Note that these transition times do not include the CIP to SSIP transition (Figure 1D in main manuscript).
